# Barbell Resolves Demultiplexing and Trimming Issues in Nanopore Data

**DOI:** 10.1101/2025.10.22.683865

**Authors:** Rick Beeloo, Ragnar Groot Koerkamp, Xiu Jia, Marian J. Broekhuizen-Stins, Lieke van IJken, Els M. Broens, Aldert Zomer, Bas E. Dutilh

**Affiliations:** Department of Biology, Science4Life, Utrecht University, Utrecht, the Netherlands; Department of Computer Science, ETH Zurich, Zurich, Switzerland; Institute of Biodiversity, Ecology, and Evolution, Faculty of Biological Sciences, Cluster of Excellence Balance of the Microverse, Friedrich Schiller University Jena, Jena, Germany; Division of Infectious Diseases and Immunology, Utrecht University, Utrecht, the Netherlands

**Author notes:** These authors contributed equally to this work (Aldert Zomer, Bas E. Dutilh).

**Keywords:** Demultiplexing, Barcoding, Sequencing, Reads, Assembly

## Abstract

**Background:** Oxford Nanopore sequencing enables long-read sequencing across diverse applications, yet the experimental artifacts introduced by Nanopore barcoding are not well characterized. These artifacts can affect demultiplexing accuracy and downstream analyses.

**Results:** We performed a rapid barcoding experiment on 66 diagnostic samples and found that 83% of reads carried the expected single-barcode pattern, while 17% contained multiple barcodes or other artifacts. Current demultiplexers, including the widely used `Dorado`, fail to correctly handle these complex cases, leaving approximately 7% of reads partially trimmed and contaminated with adapter fragments. Additional issues include the presence of two barcodes at the same read end—either identical, originating from the same sample, or different, introduced after pooling. The latter can lead to barcode bleeding when the outer barcode is incorrectly selected. To address these challenges, we developed `Barbell`, a pattern-aware demultiplexer that detects all barcode configurations. `Barbell` reduces trimming errors by three orders of magnitude, minimizes barcode bleeding, and supports custom experimental setups such as shorter barcodes, dual-end barcodes, and custom flank sequences.

**Conclusions:** Our results highlight the impact of complex barcode attachments in Nanopore sequencing and demonstrate that `Barbell` drastically reduces their effects on downstream analyses. `Barbell` is open source and available at https://github.com/rickbeeloo/barbell.

## 1 Background

Nanopore sequencing is a revolutionary technology in genomics, offering real-time, long-read DNA and RNA sequencing capabilities with minimal capital investment and laboratory footprint. Recent technological advances, particularly the introduction of the R10.4.1 pore architecture with its dual-head design and longer recognition sequence, coupled with improved basecalling models, have significantly improved sequencing accuracy to over 99% [1]. These advances allow application of Nanopore sequencing in 16S amplicon sequencing [2], genome assembly [3], and metagenomic analysis [4].

To reduce costs, multiple samples can be sequenced simultaneously through multiplexing, where unique molecular barcodes (typically 24 nucleotides) are attached to the DNA during library preparation. Barcodes are attached via tagmentation, ligation, or PCR. In tagmentation, a transposase fragments DNA and inserts barcoded adapters at the cut sites. In ligation, barcoded adapters are enzymatically joined to the ends of intact DNA fragments. In PCR barcoding, the barcode sequence is built into the 5’ end of the primers used for amplification. During PCR, these barcoded primers anneal to the target region and introduce the barcode as part of the amplified product. As a result, each amplicon carries a unique barcode corresponding to its sample, eliminating the need for a separate ligation or tagmentation step. After sequencing, software is used to detect the barcodes and assign the reads back to the original samples, called demultiplexing. Accurate demultiplexing presents significant technical challenges such as reads with multiple barcodes or poor barcode quality. While much research has focused on error rates and error correction in sequencing reads, much less attention has been given to experimental error during library preparation, although these can have serious consequences in downstream analyses such as sequence assembly or quantification.

In the late 2010s, Illumina sequencing was shown to suffer from index switching, with up to 7% of reads assigned to the wrong sample [5]. Although following experimental best practices can substantially reduce this confounder, analyses of low-abundance DNA remained at risk, such as tumour profiling [6]. Assignment to the wrong sample is described under various names, including barcode bleeding, cross-talk, and leakage.

Only a few studies have quantified barcode bleeding in Nanopore data, reporting rates ranging from 0.056% to 1.5% [7–10]. Xu et al. [8] attributed ≈80% of misassignments to concatenated reads, with the remainder due to uncertain barcodes. Wu et al. [9] argued that in *Salmonella*, where antigen-determinant loci range from 100–5000 bp, even minor barcode bleeding could alter serotype predictions. Similar concerns were raised for *Plasmodium falciparum* surveillance [10]. Thus, even small levels of barcode bleeding could compromise diagnostic accuracy.

In addition to demultiplexing, many demultiplexers also perform trimming of barcodes and adapters. However, for Illumina data this process often leaves residual adapter sequences in the reads. For instance, Moeller et al. [11] reported widespread Illumina adapter contamination in the MGnify database, particularly at contig ends.

In Nanopore data, adapter contamination has also been reported [12, 13]. Liu-Wei et al. [12] noted that untrimmed adapters often received low basecalling scores, which in turn reduced the overall read quality score. As a result, reads that were otherwise of good quality were unnecessarily discarded during quality filtering.

Overall, maximizing read-assignment rates while minimizing incorrect assignments can be method or experiment dependent and therefore remains a challenge. Hence, we argue that demultiplexers should provide extensive feedback and scores to end users to aid in understanding their data and making informed decisions.

### Barcode scoring

In Nanopore experiments the barcodes are often flanked by specific sequences, such as adapters or primers. Current demultiplexers, such as Dorado and Flexiplex, locate the flanking regions and search for barcode sequences within them. Barcodes are scored using Edlib [14], which performs a semi-global alignment based on edit distance. The edit distance, also known as Levenshtein distance, measures the number of edits required to transform one sequence into another. However, alignments with the same number of edits can still be substantially different, and do not necessarily reflect Nanopore errors [15, 16]. In RNA sequencing workflows there are promising developments that, instead of edit distance, directly use the pore signal to aid in demultiplexing [17, 18]. However, these are limited to RNA-seq set-ups and the produced models only support a subset of all barcodes. Moreover, sequencing data is almost exclusively shared as Fastq files in the sequence read archive (SRA) instead of the signal containing POD5 files. As a result, re-analysis of published data often still relies on demultiplexing from Fastq files rather than from the raw POD5 signal data.

Instead of edit distance we explore another approach, where we relate barcode scoring to general string matching problems. Specifically, we use the subsequence kernel of Lodhi et al. [19]. The idea behind subsequence kernels is that a match between two strings is better if the matching characters are close to each other. For example, **gen** requires three edits to align with “genomic” or “gnoeminc”. In the first case, the match is contiguous (**gen**omic), while in the second the matches are separated (**g**no**e**mi**n**c). Thus, under the subsequence kernel, the first alignment scores higher (see Section 5.2). Subsequence kernels are applied extensively in biological machine learning tasks [20], but here we propose to use it as additional scoring on the CIGAR representation of an edit distance alignment. The goal is to disambiguate edit distance ties, or cases where half the barcode is lost, but the remainder is sufficient to distinguish it from others.

### Custom experiments

The flexibility of Nanopore sequencing allows researchers to readily adapt Nanopore protocols, using for example different barcode configurations, primers, or other custom flanking sequences (e.g. Jia et al. [21]). `Dorado` does often not support such cases^1^. In addition, `Dorado` relies on internal edit-distance cut-offs and heuristics that do not necessarily generalize across experiments. Tools such as `Splitcode` [22] and `Flexiplex` [23] have improved flexibility by allowing users to supply their own target sequences, but important limitations remain. `Splitcode` is restricted to Hamming distance (i.e. substitutions only), so it cannot handle insertions and deletions that are common in Nanopore data [12]. `Flexiplex` does support edit distance but was developed for RNA workflows and permits only a single left-side barcode, preventing use in dual-end barcode experiments. Finally, all these tools require the user to pre-specify the pattern to search for (for example, “a single left-side barcode”). We show that in reality only ≈ 80% of the reads actually contain the expected barcode pattern, and the remainder would potentially be discarded, or could result in barcode bleeding based on users assumptions. Making users aware of these patterns by reporting them as part of the tool’s output is crucial to maximize the demultiplexing yield, and also to communicate potential experimental issues.

### Barbell

We introduce `Barbell`, an extensive tool for demultiplexing that contributes on several fronts:

- Insight into the experimental errors of Nanopore sequencing
- Overview of automatically detected barcode patterns
- Handling of complicated custom experimental set ups (e.g., multiple primers, shorter/longer barcodes, and dual-end barcodes)
- New barcode scoring scheme reflecting Nanopore errors
- User-friendly command line interface
- Presets for common Nanopore kits
- The option to only include reads displaying safe ligation/tagmentation patterns or maximize assignment (e.g., for assembly)

We compared `Barbell` with existing demultiplexers `Dorado` and `Flexiplex`. Our evaluation included trimming errors and the effects of read contamination on taxonomic assignment and genome assembly. We also assessed contamination in NCBI’s core nucleotide database to chart its broader prevalence.

## 2 Results

We developed `Barbell` to demultiplex Nanopore reads. To obtain experimental data for testing we first performed a Nanopore rapid barcoding experiment (SQK-RBK110.96) where we sequenced 66 diagnostic samples (BC01 to BC66) and a negative control (BC67). Then we explored barcode contamination in public data, and how these affect downstream analyses.

### 2.1 Demultiplexed reads

Sequencing of the 66 diagnostic samples yielded a total of 4,937,349 reads which we demultiplexed with `Dorado`, `Flexiplex`, and `Barbell`. `Dorado` assigned 4,647,221 (94.1%) to a barcode, `Flexiplex` 4,667,336 (94.5%), and `Barbell` 4,246,261 (86.0%). We note that the number of demultiplexed reads is a quantitative measure, not necessarily qualitative as we explore in the next sections. The average runtimes were 6 min 50 s for `Dorado`, 1 min 2 s for `Flexiplex`, and 5 min 50 s for `Barbell`. In Section 2.2, we introduce an alternative search pattern for ligated reads, which increased the `Barbell` runtime to 9 min 26 s. Throughout the following sections we will often refer to “patterns” as described in Section 5.3 and Section 5.4.

### 2.2 Patterns in rapid barcoding data

#### Common patterns in reads

Rapid barcoding is designed to attach a single barcode to one end of the read and we observed this pattern in 82.8% of reads (4,089,173; Table 1). In total, 709 distinct barcode attachment patterns were detected: 6.1% (299,766) of reads carried barcodes on both ends, 3.5% (173,692) contained two barcodes on the left, and 1.0% (46,707) carried both two left barcodes and a single right-end barcode. Although rare, some reads consisted almost entirely of barcodes, with up to eight in a single read (Figure S3; Additional file 1). Overall, ≈17% of reads deviated from the expected design.

**Table 1:**
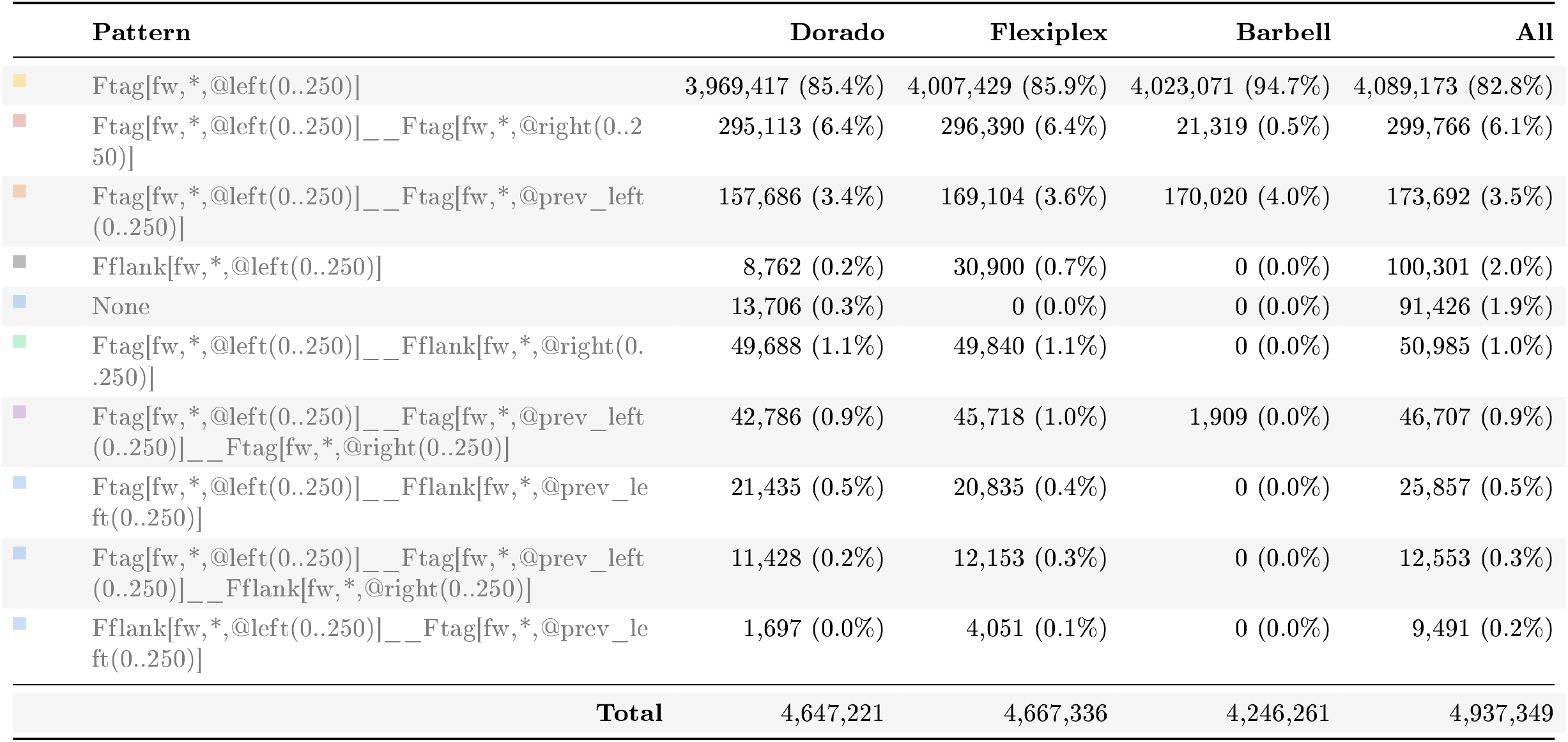
Top 10 barbell patterns in rapid barcoding. This table shows the 10 most common out of 709 total patterns detected in the reads. The “Color” column contains the color used in the plots throughout this manuscript. The “Pattern” column shows the read pattern assigned by `Barbell` (see Section 5.3). The “All” column is based on the `Barbell` `annotate` output, and the other columns show the number of reads with the “Pattern” that were trimmed and output by `Dorado`. `Flexiplex`, and `Barbell`. The percentages are based on the total number of demultiplexed reads by each tool (Total). The discrepancy between the “All” count and the count, in the “Barbell” column corresponds to the number of cases where the full read sequence was trimmed, e.g. when the entire read consisted of barcodes or flanks. The None pattern indicates that `Barbell` did not find any barcodes and thus no pattern was assigned to those reads.

#### Incorrectly trimmed reads

To detect contamination in trimmed reads, we searched all trimmed reads for flanks and barcodes using `Sassy` [24] based on edit distance (see Methods). Among the demultiplexed and trimmed reads, Nanopore adapter and barcode remnants (hereafter “contamination”) were detected in 10.0% of reads trimmed by `Dorado` (n=464,518), 8.8% by `Flexiplex` (406,450), and 0.004% by `Barbell` (166). The few remaining contaminanted reads detected after `Barbell` trimming can be explained by the prefix-based search mechanism of `Sassy`, which `Barbell` itself also uses to locate barcodes and flanks. Because `Sassy` assigns a lower cost to missing prefixes—allowing for partially truncated barcodes near read ends—secondary barcodes in double-barcoded reads may only become detectable after removal of the primary prefix.

Especially short reads (≤ 250 bp; 880,637 in total) were not consistently trimmed across tools. `Dorado` retained 88.1% (775,409) of short reads after trimming, `Flexiplex` 89.0% (783,496), and `Barbell` 43.2% (380,308). Here, “retained” means that the reads were not completely trimmed away — in other words, they were not composed entirely of barcode sequence according to the tool. Among the retained reads, remaining contamination was detected in 44.5% (345,142) of those trimmed by `Dorado`, 40.3% (315,877) by `Flexiplex`, and only 0.04% (160) by `Barbell`. In `Dorado` and `Flexiplex`, contamination was primarily associated with reads carrying multiple barcodes—either two left barcodes or a barcode at both ends (Figure 1A).

**Fig. 1:**
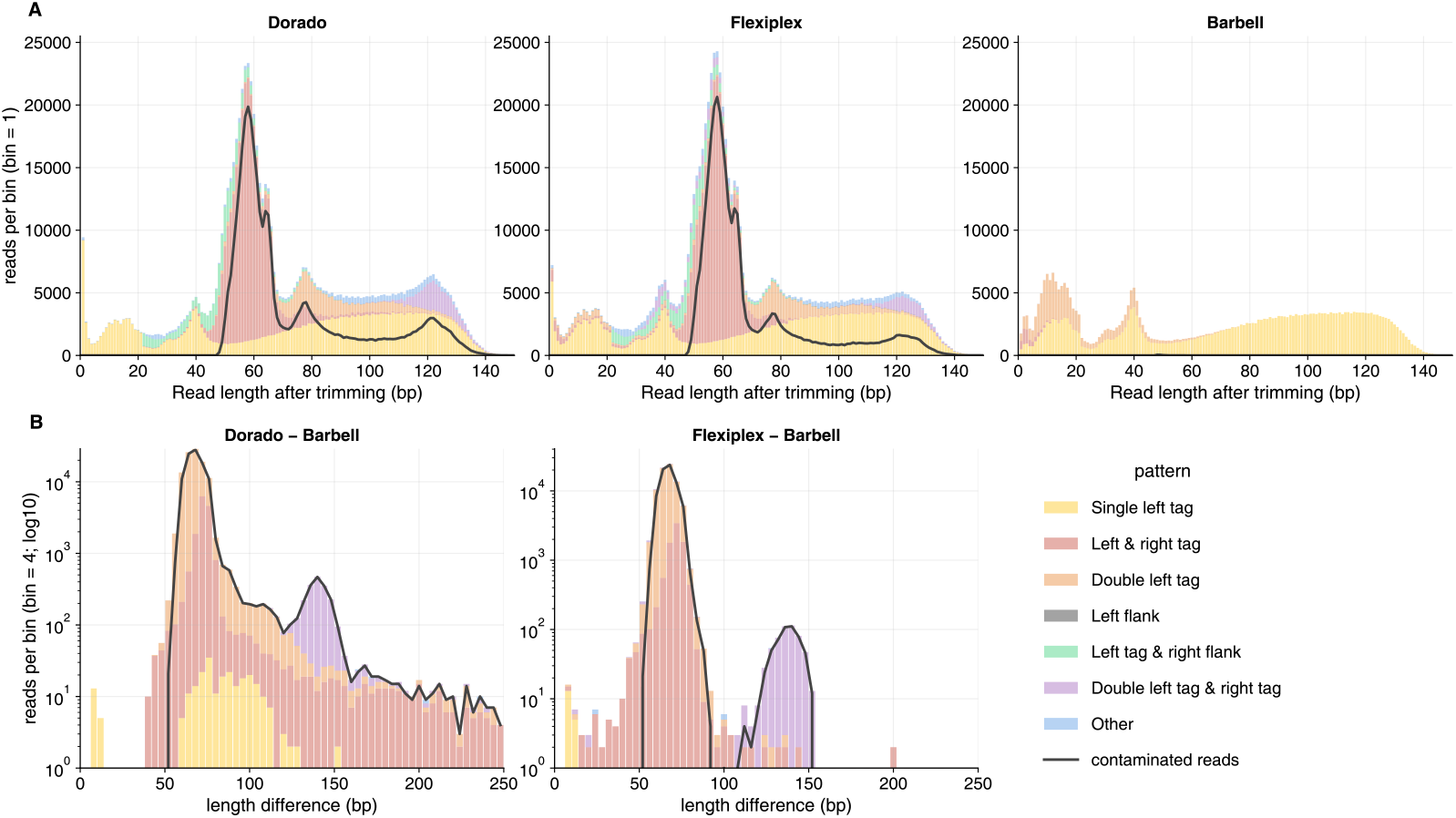
Trimmed read length comparisons. **(A)** Distribution of read lengths after trimming short reads (≤250 bp) for `Dorado`, `Flexiplex`, and `Barbell`. Bars are colored according to the pattern assigned by `Barbell`, as outlined in Table 1. Most reads were trimmed from 250 bp to ≤150 bp (280 reads >150 bp not shown). `Dorado` and `Flexiplex` produced many trimmed reads of ≈60 bp, visible as a prominent peak, which were absent in `Barbell` output. These ≈60 bp trimmed reads originated from sequences containing two adjacent barcodes without sequence inbetween: `Dorado` and `Flexiplex` recognized only one barcode and output the remaining barcode sequence as a valid read, whereas `Barbell` detected both barcodes and removed the entire read as contamination. The black line indicates trimmed reads that contained detectable Nanopore adapter sequences (see Methods), which closely tracked the ≈60 bp contamination peak, confirming these were artifact sequences rather than genuine biological reads. **(B)** Trimmed read length differences for input reads >250 bp comparing reads trimmed by `Dorado` vs. `Barbell` (left) and `Flexiplex` vs. `Barbell` (right). Note the logarithmic y-axis. Both `Dorado` and `Flexiplex` output longer trimmed reads than `Barbell`, often corresponding to a single undetected (≈60 bp) or two undetected (≈120 bp) barcodes, similar to those in **(A)**. As the difference between the tools was generally ≤10 bp. we only showed differences >10 bp.

Because `Dorado` and `Flexiplex` trimmed only one of the barcodes, additional copies remained, producing characteristic peaks: one remaining barcode resulted in a peak at ≈ 60 bp, and two remaining barcodes resulted in a peak at ≈ 120 bp. If complete rapid barcoding sequences would remain, peaks would be expected at multiples of 90 bp, corresponding to the full rapid barcoding sequence. However, as discussed in Section 2.2, having two adjacent barcodes in a read was often paired with the loss of ≈ 30 bp that shifted these expected lengths.

Also longer reads (>250 bp) were not trimmed consistently by the different tools. 423,908 reads trimmed by `Dorado` and 402,780 reads trimmed by `Flexiplex` were longer than those trimmed by `Barbell`. The length difference was generally small, but for 107,890 `Dorado` and 80,496 `Flexiplex` reads the difference exceeded 10 bps, of which 93.1% (100,496) and 91.7% (73,817) contained contamination, respectively (Figure 1B). For genome assembly, typically only trimmed reads ≥ 1,000 bp are used. Also in these longer reads contamination was observed in 59,563 `Dorado` reads, 43,656 `Flexiplex` reads, and 3 `Barbell` reads.

Overall, incorrect trimming affected >8% of reads when using `Dorado` and `Flexiplex`. Most contamination was seen for short reads, however persisted in reads exceeding 1000 bp.

#### Double barcode attachment and bleeding

Double left-end barcodes were identified in 173,692 reads (3.5% of total; Table 1). In 30.7% of these reads (53,386), the right flank of the first barcode was directly “fused” to the second barcode, resulting in complete loss of the left-flank sequence and frequent partial deletion of the second barcode (Figure 2). Consequently, the mean edit distance to the first barcode was 3, compared to 7 for the second. We observed fusions for all barcodes, but the prevalence of fusion-associated deletions in the first 6 bp of the second barcode varied by barcode, for example: BC05, 95.4% (2,073/2,173); BC25, 95.8% (2,106/2,198); BC61, 29.5% (901/3,053); and BC45, 52.3% (2,027/3,874). We observed similar patterns when analyzing public datasets (Weinmaier et al. [25]: BC05, 372/393, 94.7%; Di Pilato et al. [26]: BC45, 61/202, 30.2%). Scanning all untrimmed reads for the fusion pattern revealed that 3.3% of all reads (n=165,396) contained such a double-barcode fusion. We hypothesized that some sequence at the fusion points might remain uncalled by the basecaller, producing detectable pore signals without corresponding basecalled bases. To investigate this, we examined the raw signals at these sites (see Section B; Additional file 1), but did not observe any systematic deviations. Nevertheless, these fusions thus shows a characteristic loss of sequence that complicate detection of the second barcode.

**Fig. 2:**
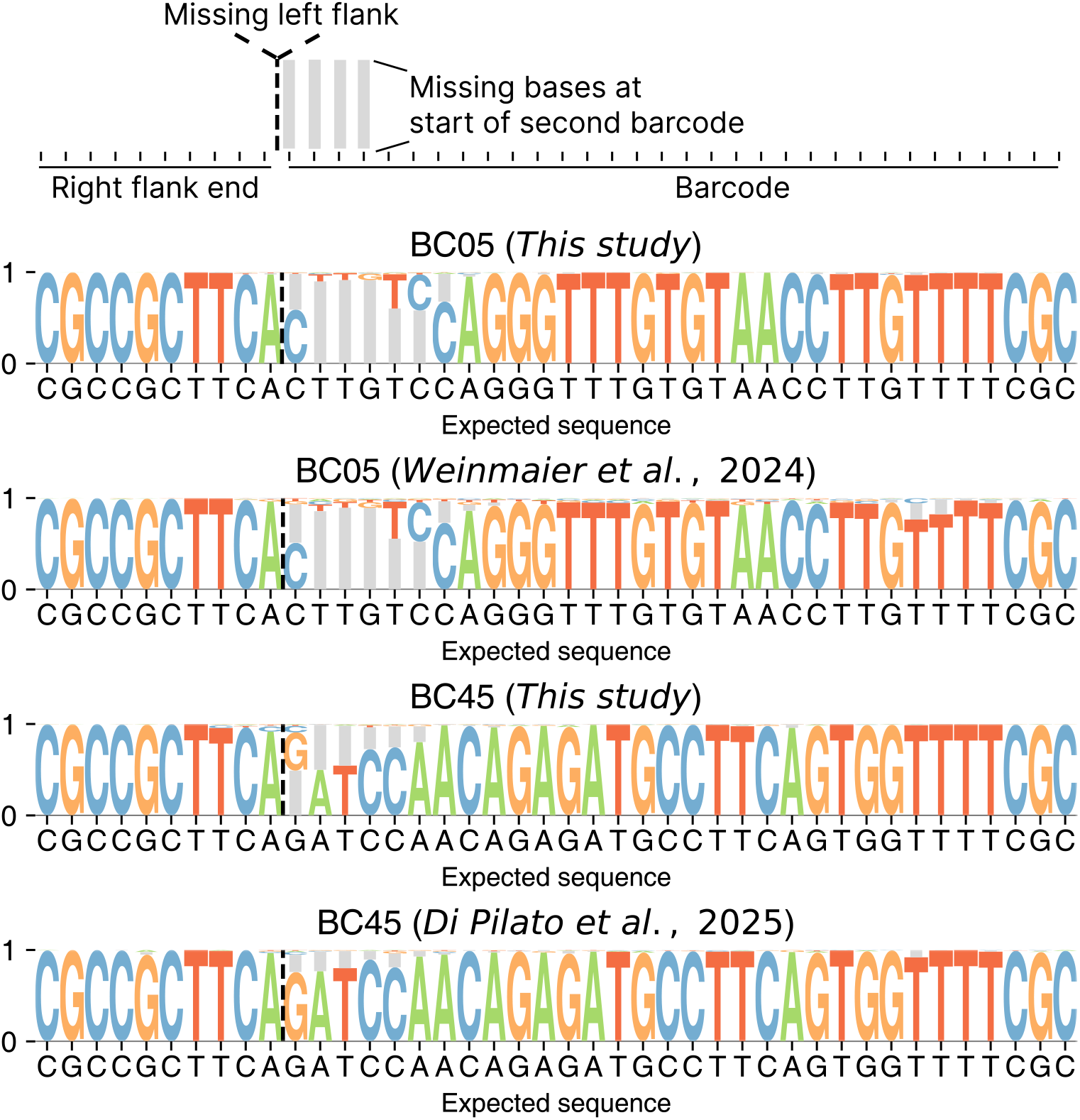
Detection of fused rapid barcodes and associated deletions. Among 173,692 reads with two left barcodes, 53,386 showed an unusual arrangement in which the right flank adjacent to the first barcode was directly fused to the second barcode (see Section 5.7). The sequence logos show the fusion junctions for BC05 and BC45 in our and public datasets. Letter height indicates base frequency; gray bars mark alignment gaps (missing bases). In typical reads, a left flank, barcode, and right flank are observed in order, whereas fusion reads show the right flank of the first barcode (ending with TTCA, dashed line) joined directly to the second barcode, always lacking its left flank (not shown) and first part of the second barcode. Deletions within the first 6 bp of the second barcode occurred in 95.4% (BC05) and 52.3% (BC45) of our reads, and with comparable frequencies in public datasets (94.7% and 30.2%, respectively). Loss of the first 1–6 bp of the second barcode was thus more frequent in fusions involving BC05 than BC45.

Failing to detect the second barcode would be problematic when the two barcodes were different. Of the 173,692 reads carrying two left barcodes, 99.5% (172,759) contained the same barcode twice. While such duplications impaired `Dorado`’s trimming (Fig. 1), it did not affect the demultiplexing. In 933 reads (0.5%), however, the tools disagreed: `Dorado` consistently reported the outer copy, whereas `Barbell`, which detects both instances, assigned the read to the inner barcode. To evaluate which assignment was correct, we compared read-level taxonomic annotations with those of the assemblies linked to the assigned barcodes. This approach is limited by the uncertainty of read-level annotations (here filtered at ≥ 100 bp hit length; see Section 2.2) and by the fact that nine species were present in two samples (Table S1; Additional file 1). Taxonomic annotation supported the inner barcode in 513 cases (55.0%) versus 46 (4.9%) for the outer, with the remainder being unclassified.

These results indicate that barcode misassignment in `Dorado` arose from the selection of the outer copy or failure to detect the inner copy (Figure 1). `Barbell` identified the inner barcode and was thus less affected by such experimental artifacts.

#### Incorrect trimming and taxonomic assignment

Next, we evaluated how contamination affected the taxonomic assignment of reads. The reads trimmed by `Dorado` and `Barbell` were annotated using `Centrifuger`, which assigns taxonomy based on kmer matches between the reads and a reference database (RefSeq [27] here). If a read has matches to multiple taxonomically different entries, `Centrifuger` moves up the taxonomic hierarchy and reports the lowest shared taxonomic rank across all matches.

In total, 4,729,126 reads received a taxonomic assignment. For each read, we compared the classification obtained after trimming with the two different tools. Identical taxonomic assignments were obtained for 3,882,881 reads (82.1%), while 392,499 reads (8.3%) differed, either being unclassified by one tool or assigned to different families. Most discrepancies originated from reads trimmed by `Barbell` that remained unclassified at the family level, whereas the corresponding `Dorado`-trimmed reads were assigned to the *Enterobacteriaceae* (n=203,106; 51.5%), predominantly *E. coli* (n=39,967; 10.2%). Among the *Enterobacteriaceae* assignments, 162,044 (41.3%) required hierarchical resolution, and `Centrifuger` therefore did not associate a specific RefSeq accession with the taxonomic assignment. For the remaining 26,759 reads, the assignments were based on 239 unique RefSeq entries. To investigate the source of ambiguous *Enterobacteriaceae* assignments among trimmed `Dorado` reads, we examined whether residual *Mu*-transposon sequences from the Nanopore Rapid Barcode flanks might have matched endogenous *Mu* transposons in these bacteria. The 26,759 `Dorado`-trimmed reads were aligned to the 239 RefSeq genomes, and the genomic regions within 5 kb of the alignment sites were analyzed. Most alignments (24,158 reads; 90.3%) were located near genes characteristic of transposons, such as those encoding a *recombinase family protein*. Notably, many of these regions also contained phage-associated genes, including those encoding the *tail fiber assembly protein* (24,159 reads; 90.3%) and the *Mu* phage-specific *Mom* family adenine-methylcarbamoylation protein (24,157 reads; 90.3%). The consistent co-occurrence of transposon- and phage-related genes strongly indicates the presence of *Mu* phage integration sites. Alignment of the *Mu* phage genome further confirmed this, with 90.3% (n=24,161) of reads mapping within *Mu* phage regions. Thus, residual *Mu* transposon sequences from tagmentation, when untrimmed, created artificial matches to endogenous *Mu*-like elements in reference genomes, leading to misleading taxonomic assignments.

The Genome Taxonomy Database (GTDB) is often used for taxonomic annotation as its high quality sequences are expected to yield accurate assignments. Using `Centrifuger` with the GTDB resulted in 343,285 family-level discrepancies between `Dorado`-trimmed and `Barbell`-trimmed reads. Where `Barbell`’s reads were unassigned, `Dorado`’s reads were mostly assigned to *Balneolaceae* (n=101,443; 29.6%) and *Streptomycetaceae* (n=95,341; 27.8%). We traced these matches back to contamination in public assemblies (see Section 2.4). Specifically, 67.4% of *Streptomycetaceae*, all *Streptomyces* species, were assembled by Jørgensen et al. [28]. All *Balneolaceae* were from a single *Gracilimonas* assembly (`GCF_040117685.1`) by Lim et al. [29].

Because the rapid barcoding region spans only 90 bp (see Methods), we suspected that limiting taxonomic assignments to matches ≥ 100 bp would reduce the effect of rapid barcoding contaminants. This was indeed the case, lowering discrepancies to 130 reads at the family level, but also reducing the total number of assigned reads by 25.3% (4,169,566 to 3,131,098).

#### Propagation into assemblies

Incomplete trimming also impacted genome assemblies. Assemblies were successfully generated for 64 of 66 samples; BC02 and BC03 contained too few reads for assembly. Of the 64 assemblies, 59 were bacterial and 5 fungal (Table S1; Additional file 1). For the bacterial assemblies, CheckM2 [30] estimated completeness and contamination at 99.42% and 1.13% for `Dorado`, and 99.26% and 1.03% for `Barbell`, respectively. However, tools like CheckM2 evaluate contamination based on single-copy marker genes, and these values do not directly reflect the presence of residual artificial sequences. A straightforward approach to detect experimental contamination is to screen all 64 assemblies for residual rapid barcoding flanks and barcodes. This analysis revealed contamination in seven assemblies from `Dorado` trimmed reads (BC19 (3×), BC21 (1×), BC35 (6×), BC39 (1×), BC49 (3×), BC58 (1×), and BC64 (1×)), whereas no contamination was detected in assemblies generated with `Barbell` trimmed reads.

In the *Saccharomyces cerevisiae* assembly for BC49, we identified contamination at three locations. One at the start of a 23,203 bp contig (positions 1 to 92) that originated from a double-left barcode read that `Dorado` failed to trim. This residual sequence extended the contig, with additional contaminated reads mapping to it (Figure 3). The same read was correctly trimmed by `Barbell`, preventing contamination of the assembly. BLAST analysis of this contig showed a near-perfect alignment to *S. cerevisiae*, except for the first 86 bp, which instead matched diverse taxa including *Pseudomonas aeruginosa, Photobacterium leiognathi*, other bacteria, and synthetic constructs. Thus, the first ≈9O bp of the contig are indeed generally absent from *S. cerevisiae* genome sequences and instead matched contamination or endogenous *Mu* transposons in public databases (later in Section 2.4).

**Fig. 3:**
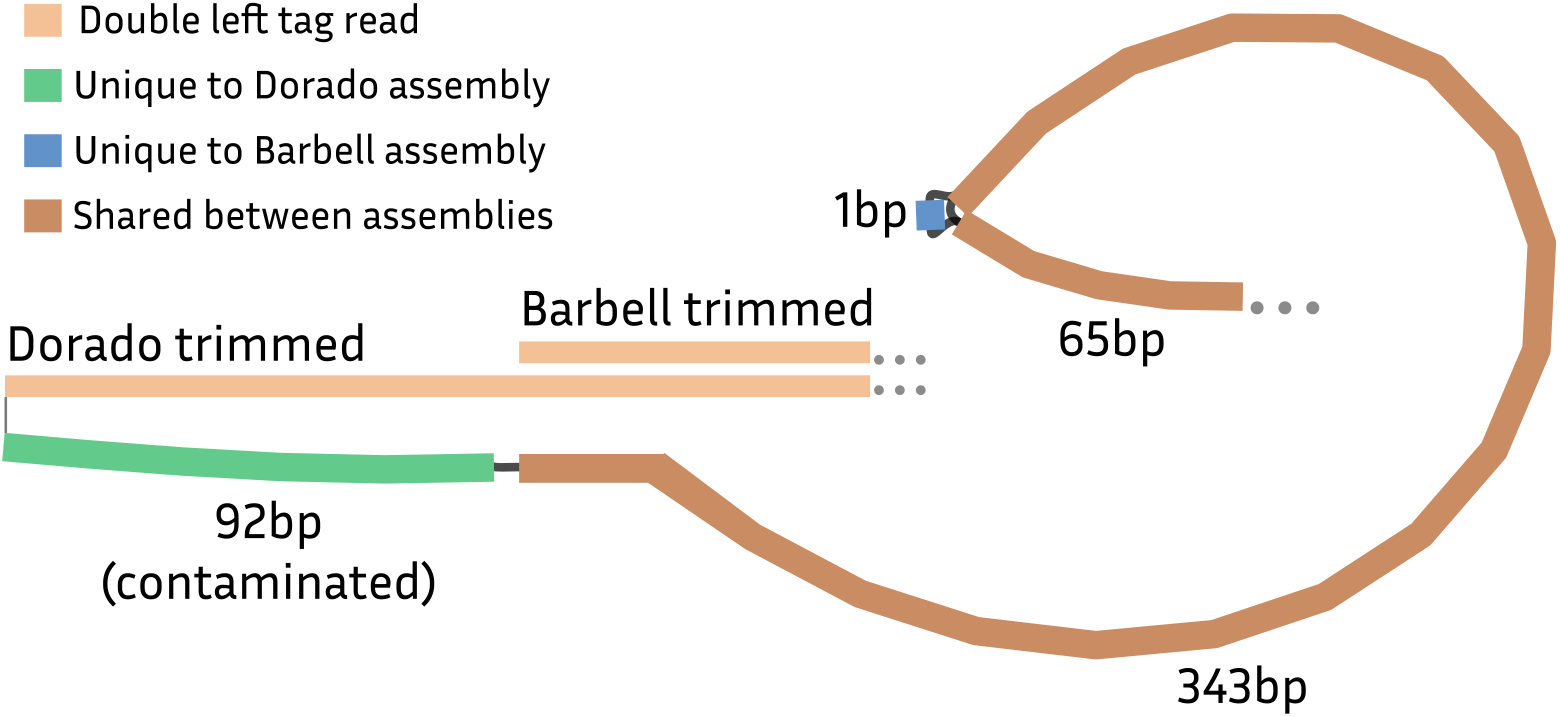
Merged assembly graphs for a *Saccharomyces cerevisiae* contig. Genome assembly of a single contig from *Saccharomyces cerevisiae* from `Dorado`- and `Barbell`-trimmed reads (23,203 bp vs. 23,050 bp). Shown are the first 500 bp of the `Dorado` assembly and the corresponding region from the `Barbell` assembly. The nodes represent unitigs, and the edges their connections. The assemblies were identical except for the first 92 bp and a single nucleotide difference. The extra 92 bp in the `Dorado` assembly originated from a single barcode sequence left untrimmed by `Dorado`. The difference was caused by one read containing two left barcodes; `Dorado` removed only the outer barcode, leaving the inner one intact which got incorporated in the assembly. In contrast, `Barbell` removed both barcodes, preventing this contamination.

Overall, `Dorado` frequently failed to trim reads containing multiple barcodes, leaving residual sequences that affected both taxonomic annotation and assembly. In contrast, `Barbell` effectively removed such experimental artifacts, mitigating their downstream impact.

### 2.3 Comparing scoring schemes

Overall, `Dorado` demultiplexed 459,987 more reads than `Barbell`. Most of these (92%, 428,823) corresponded to reads that `Barbell` had annotated but excluded from its final output, either because they did not match rapid-barcoding patterns or because trimming produced empty sequences.

To assess whether these additional reads were correctly assigned, we compared species-level taxonomic annotations of the trimmed reads with those of the assemblies linked to their assigned barcodes. Only 6.8% (31,164) of `Dorado`’s additional reads showed consistent taxonomic assignments, indicating limited accuracy among these extra demultiplexed reads.

A total of 13,706 reads were missed by `Barbell` because their rapid-barcoding flanks exceeded the automatic cutoff of 20 edits (Table 1); `Dorado` correctly demultiplexed 72.0% (9,857) of these. Another 8,762 reads were annotated as Fflank by `Barbell` when subsequence scoring was inconclusive, 71.3% (6,244) of which showed the expected taxonomy. Such cases can be recovered by lowering `Barbell`’s subsequence-scoring thresholds (Section 5.2, Section 5.5).

Conversely, `Barbell` demultiplexed 81,931 reads that `Dorado` failed to assign. For 69.3% (56,798) of these reads, the species assignments were consistent with the corresponding assemblies, with the remainder unclassified.

As illustrated in Figure 2, barcode fusions frequently resulted in partial loss of the second barcode. Such events are difficult to detect using simple edit-distance scoring, as the missing initial bases increase the apparent distance by roughly four edits. Because `Dorado` detects only the first barcode, direct comparison of scoring between the two tools is not possible. `Dorado` requires a minimum difference of three edits between the two best matches, which would often prevent assignment of truncated barcodes (Figure 4). In contrast, `Barbell`’s subsequence-based scoring successfully identified these cases.

**Fig. 4:**
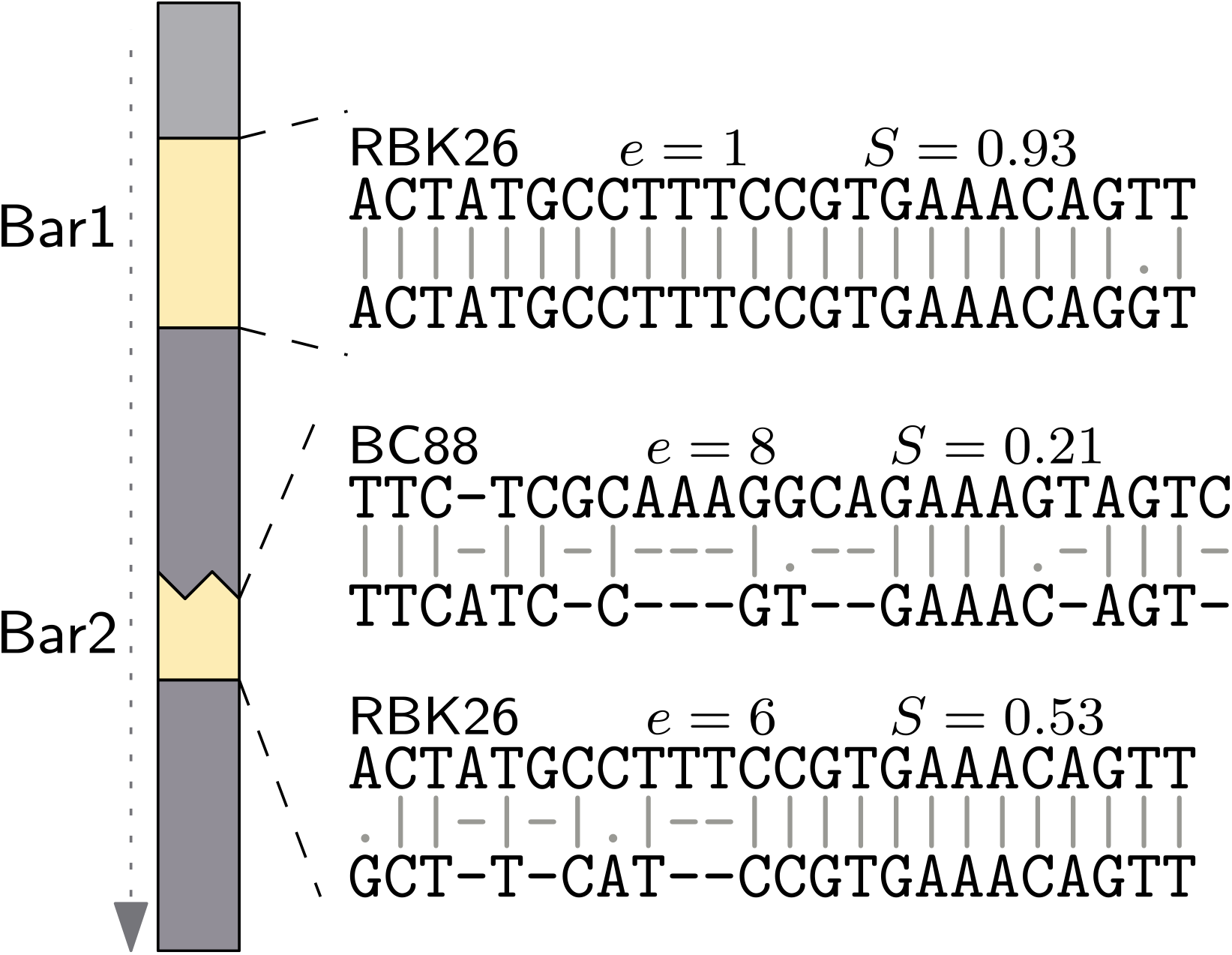
Example of scoring in a double-left barcode read. Example of a read containing two left barcodes (Figure 2), where the second barcode (Bar2) is truncated. The first barcode (Bar1) is unambiguously assigned to RBK26 with a single edit. For the second barcode, edit distance alone yields two close matches; to RBK26 and BC88 (6 and 8 edits, respectively), a difference too small to be assigned by `Dorado`. Subsequence scoring strongly favors RBK26, as 12 of 24 consecutive nucleotide matches provide enough evidence for RBK26 over the more interleaved error pattern for BC88. This example highlights how partial yet contiguous matches can enable barcode recovery, even when edit distance alone might not be discriminative.

In summary, `Barbell` recovered 56,798 reads that `Dorado` failed to assign, while missing 19,950 reads that `Dorado` likely demultiplexed correctly. Overall, `Barbell` provided substantially cleaner trimming and more robust handling of complex barcode patterns (Section 2.2).

### 2.4 Barcodes and their flanks in public databases

Adapter contamination in genome assemblies has previously been reported for Illumina data [11]. Because we observed that Nanopore sequences are not always removed by standard tools such as `Dorado`, we queried the NCBI “core nucleotide” database (≈ 810 GB) for Nanopore contamination.

#### Match statistics

The search identified 103 matches to both rapid barcode flanks and barcodes, including 68 exact matches (0 edits) across 67 assemblies. Additional hits were found to the flanking sequences alone, without the barcode; these were excluded, as they may represent matches to endogenous *Mu* transposons (Section 2.2). For native flanks and barcodes, we detected 462 matches, of which 270 were exact matches across 284 assemblies. All match tables were uploaded to Zenodo^2^

#### Rapid barcodes

The most striking case was *Photobacterium leiognathi* strain SV5.1 (`CP131573.1`) where we detected BC86 11× across a 1.43 Mb contig. Unlike in our assemblies, barcodes were scattered internally, reflecting scaffolding of smaller contigs separated by N stretches. Based on the supplementary data of the corresponding paper ([31]), their initial assembly contained over 30 contigs, and scaffolding was used to reduce this to 2 contigs [31]. Re-downloading the 89,663 raw reads from the SRA database revealed that 73,580 (82%) contained the expected single-flank pattern Ftag[fw,*,@left(0..250)], 1,299 reads (1.4%) contained a double-flank arrangement Ftag[fw,*,@left(0..250)]_Ftag[fw,*,@prev_left(0..250)], followed by the same patterns as observed in our rapid barcoding experiment (Table 1). Trimming the reads with `Barbell` using the default options for the SQK-RBK114-96 kit followed by assembly produced three circular contigs (3,176,913 bp; 1,497,394 bp; 15,997 bp) and a small linear contig (4,109 bp). This is in line with the chromosomal arrangement of *P. leiognathi* [31]. The 1.49 Mb contig matched `CP131573.1`, without barcodes, showing that careful read-level trimming can improve the assembly.

Another example was an 39,350 bp *E. coli* plasmid (`CP165501.1`) [32], with BC10 contamination on the left (positions 25-111) and BC09 contamination on the right (39,256-39,329) of the contig. Notably, BC10 was present in the forward orientation, and BC09 in reverse complement. We downloaded all 113,399 raw reads from the SRA database and demultiplexed these using `Barbell` (default; SQK-RBK114-96 kit). Of the reads, 97,461 (86%) contained the expected single-flank pattern followed by those in Table 1. While 76.4% of the reads were assigned to BC09 by `Barbell`, 13.1% of the reads contained BC10 according to `Barbell`. Assembling the by `Barbell` trimmed reads for just BC09 produced a 43,165 bp contig matching `CP165501.1` from positions 111 to 39,256 corresponding to the removal of the barcode contamination of both sides of the original uploaded sequence. Thus, we identified two distinct barcodes in `CP131573.1` resulting from erroneous demultiplexing, and evidence of assembly contamination likely caused by incomplete trimming. Other cases included plasmids, mobile elements, and assemblies from Jørgensen et al. [28], *Streptococcus thermophilus* (`CP072431.1`), and *Staphylococcus aureus* (`CP150769`).

#### Native barcodes

Contamination from native kits was more widespread, spanning viruses, bacteriophages, bacteria, parasites, fungi, short rRNA sequences (≤ 1.5 kb), and organellar genomes. Most reminant native sequences were detected in *Mycolicibacterium novocastrense* (`CP097264.1)` with 50 matches to NB02 [33]. Other examples include human SARS-CoV-2 (`OV192362.1`), the house cricket densovirus (`PP054203.1`), bacteriophages (`OP583592`, `OR487170.1`, `PP989835.1`), mitochondrial DNA from *Tonna galea* (`NC_082277`), and chloroplast DNA from *Cephaleuros karstenii* (`NC_060534`).

An illustrative plasmid case was `CP142556.1`, an 8,423 bp *E. coli* ExPEC_A376 plasmid [34]. Although annotated as circular, a reminant NB13 was detected at positions 8376–8421. Self-alignment revealed an overlap from bases 1–35 to 8341–8375, leaving the barcode as an overhang. Subsequent Illumina polishing by the authors did not remove this artifact, showing that circularity calls alone cannot guarantee contamination-free sequences.

Thus, both rapid and native Nanopore barcoding kits have left detectable footprints in public databases across viruses, bacteria, plasmids, organelles, and rRNA records.

### 2.5 Barbell tool: usage and applications in custom experiments

Untrimmed Nanopore barcodes were common in both our datasets and public assemblies. Moreover, failing to detect multiple barcodes could lead to barcode bleeding. To address these issues, we developed `Barbell`, a Rust-based tool for accurate barcode detection, trimming, and pattern analysis (https://github.com/rickbeeloo/barbell).

Barbell increases detection accuracy and drastically reduced trimming errors. For standard Nanopore kits (for example, SQK-RBK114-96), a single command automatically identifies flanks and barcodes, sets cut-offs, performs trimming, and generates summary statistics (Figure 5). The tool further accommodates custom experimental designs, including dual-end barcodes and mixed amplicon datasets, by allowing users to define their own flanking sequences and barcodes.

**Fig. 5:**
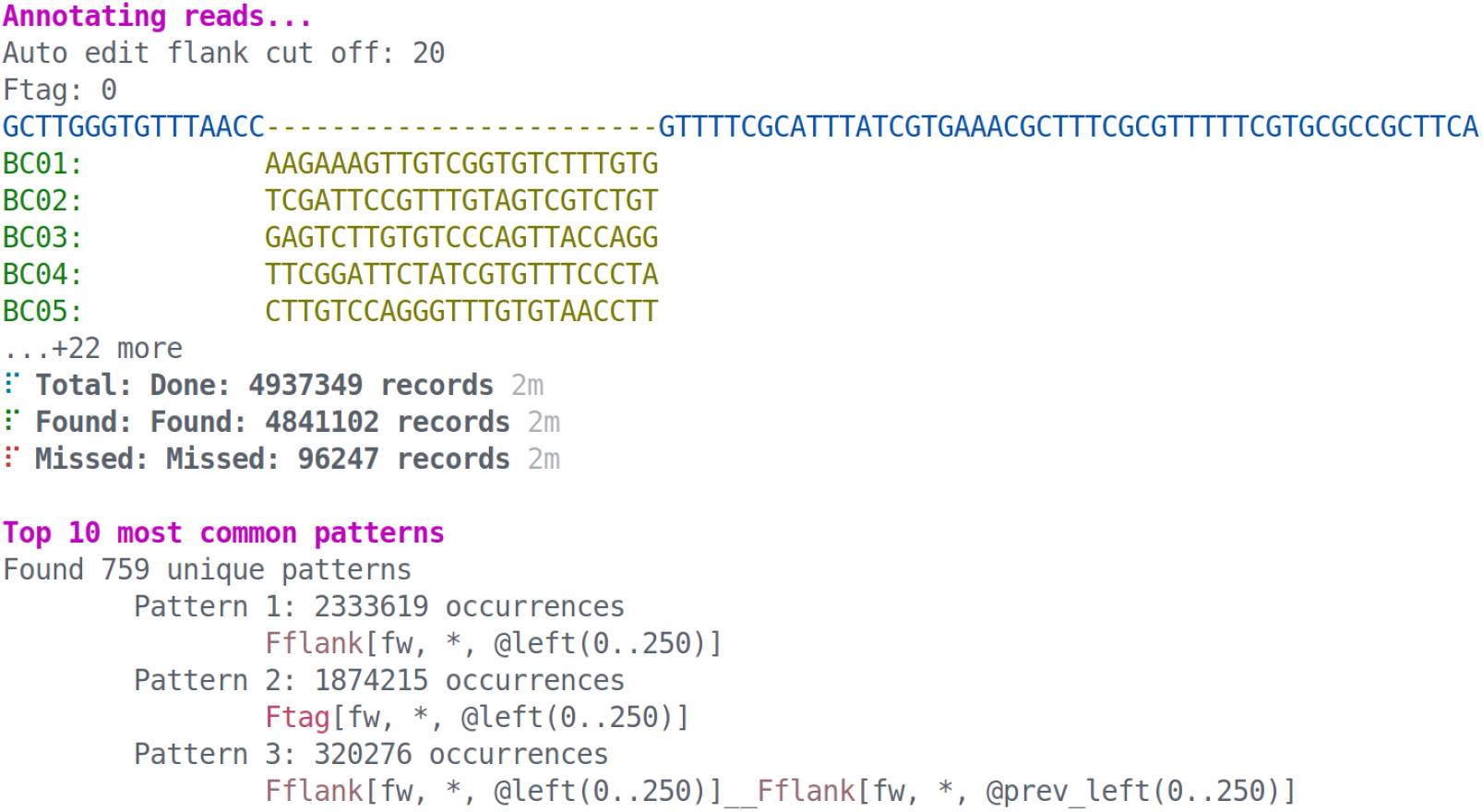
Barbell command-line interface. Example output when running `barbell kit -kit SQK-RBK114-24 -i reads.fastq -o output` The interface displays kit information, including whether the `--maximize` option was used (see Methods). Inferred flanks (blue), detected barcodes (yellow), and the automatically assigned flank edit-distance cutoff (20 in this example) are shown. The output reports progress at each step and summarizes the most frequent sequence patterns in the input FASTQ file (3/10 shown), providing a direct overview of double attachments and other experimental artefacts.

For example, Jia et al. [21] required orientation-aware demultiplexing, in which the combination of barcodes BC01–BC02 in forward–reverse orientation represented a different sample than in reverse–forward (BC02-BC01).

## 3 Discussion

We showed that `Barbell` is a powerful demultiplexing tool that provides insight into the adapter and barcode patterns in Nanopore reads. It substantially reduced trimming errors and minimized barcode bleeding. Ultimatively producing cleaner reads for downstream analysis.

### Barcode patterns

A large portion of research on Nanopore sequencing focuses on establishing error rates and mitigating their effects in downstream analysis by generating consensus sequences [35–37]. However, much less attention has been paid to what happens to reads during the experimental steps such as tagmentation and ligation. We show that only about ≈80% of reads in rapid barcoding experiments are of the expected configuration, while the remaining ≈20% contain multiple barcodes in different configurations.

Specifically reads having two barcodes on the left, or barcodes on both ends of the read are problematic for existing tools (Figure 1). We showed that in case of double-left barcodes the second copy often lacks the entire left flank and a partial prefix of the second barcode (Figure 2). As to the physicochemical mechanism involved, we did not observe any abnormal spikes in the pore signal that could indicate secondary structure. This suggests that these sequences are single stranded. There has been a recent report of biases in Nanopore sequencing related to the *mu* target site [38] which might play a role. Nevertheless, such fusions are difficult to demultiplex for two reasons. First, entire loss of the left flank makes it hard to locate the barcode region in the first place, and second, prefix loss of the barcode increases its edit distance to the reference and lowers it to other barcodes. We specifically added this fusion pattern to `Barbell` (enabled by `--use-extended`) and showed that the subsequence scoring is robust to losing a prefix (Figure 4).

While identical double barcodes on the left side heavily impaired trimming of `Dorado`, around ≈0.5% of the double left reads had two different barcodes. `Dorado`’s selection of the outer barcode here is a source of barcode bleeding.

### Contamination

Contamination from reagents and kits is well known to affect downstream analyses [39]. Many metagenomes in the MGnify database contain Illumina adapter contamination [11], and we demonstrated that also Nanopore barcodes and their flanking sequences can appear in assemblies when using existing demultiplexers. Moreover, our analysis of the “core nucleotide” database suggests that many public assemblies contain Nanopore contamination. A valuable next analysis, would be to repeat such analyses for other public resources such as GTDB which is commonly used for taxonomic assignment. The presence of such contamination can generate spurious taxonomic signals, particularly when untrimmed barcodes and flanks from existing demultiplexers form misleading links with contamination in the public databases (Section 2.2). Therefore, researchers should exercise caution when using public data as reference material. Special attention should be paid to whether matches stem from barcodes or flanking regions. Stricter post-processing rules—for example, requiring matches of at least 100 bp—can help reduce such spurious matches.

We also note that tools like Porechop [13] which is currently unsupported and Fastp-long [40] have likely played an important role in mitigating contamination by removing remnant adapter sequences after running other demultiplexers. We showed that `Barbell` can already address this issue at the initial read-processing stage, which is particularly beneficial in cases involving double-ligated barcodes where the inner barcode should be chosen.

### Usability

Researchers are increasingly developing custom experiments using their own barcodes, primers, or other tags, often resorting to custom demultiplexing scripts. `Barbell` is specifically designed for such cases as a modular tool, where the flank and barcode sequences can be changed to fit the user’s needs. Moreover, `Barbell` is currently the only tool that provides a comprehensive overview of patterns in the data. This functionality supports experimental design, helps detect potential issues, and enables subsetting of reads based on expected patterns.

## 4 Conclusion

We demonstrated that commonly used demultiplexers leave approximately 10% of Nanopore reads improperly trimmed, which can significantly impact downstream analyses such as taxonomic annotation and genome assembly. These effects are further exacerbated by the presence of similar contamination in public databases, including the core nucleotide database and GTDB, which can create artificial connections lacking true biological meaning. To address this issue, we developed `Barbell`, a pattern-aware demultiplexing tool capable of detecting complex barcode attachment patterns. `Barbell` reduced barcode bleeding and trimming errors by three orders of magnitude, demonstrating its robustness as a demultiplexer for sequence analysis.

## 5 Methods

### 5.1 Problem definition

Given a set of reads (typical length ≈10-30kb) and a set of tags (≤250 bp) determine for each read the location of the tags, and extract the trimmed reads, that is, the part of the read flanked by one or more tags. Here tags are barcodes and their flanking sequences can be adapters or other sequences such as primers.

### 5.2 Preliminaries

In this manuscirpt, we address the problem of demultiplexing, where the goal is to locate a *tag*, denoted by *τ*, of length |*τ*|, within a read *R* − *r*_0_…*r*_*n*−1_ of length *n*:*=* |*R*|. Both *τ* and *R* are strings over the DNA alphabet Σ = {*A, C,G,T*} extended with IUPAC ambiguity codes (e.g. `N, R, Y, M`). Let *σ*:= |Σ| denote the alphabet size.

Each tag *τ* consists of three parts (or substrings): a left flank *F*_*ℓ*_, a barcode *B*, and a right flank *F*_*r*_. We denote their respective lengths as |*F*_*ℓ*_|, |*B*|, and |*F*_*r*_|, such that *τ = F*_*ℓ*_ ∘ *B* ∘ *F*_*r*_, where ∘ denotes string concatenation. The barcode *B*, typically 24 bp, comes from a set of *g* known barcodes, *β =* {*b*_1_, *b*_2_, …, *b*_*g*_}, whereas *F*_*ℓ*_ and *F*_*r*_ are fixed strings with lengths varying based on the protocol, from |*F*_*r*_| = 8 for native barcoding kits, to for example |*F*_*r*_| = 50 for rapid barcoding.

Since all the barcodes share the same flanks, we can speed up searching by first locating *F*_*ℓ*_ and *F*_*r*_ and searching the barcode between them. We do this by replacing *B* in *τ* by a wildcard region, or “mask” which consists of N characters, each of which can match any character in Σ. We denote a mask of length *s* as *N*_*s*_, and use *τ*_*N*_ to represent the tag with the mask, *τ*_*N*_*:= F*_*ℓ*_ ∘ *N*_|*B*|_ ∘ *F*_*r*_. We write *R*[*i*… *j*]:*= r*_*i*_… *r*_*j*−1_ to denote a right-exclusive substring of *R* (i.e. [*i, j*)).

Throughout the manuscript we use two ways of penalizing/scoring sequences. The first measure is the edit distance. The second is a subsequence-based scoring function. We use edit distance to locate *τ*_*N*_ in the read. Subsequence scoring is used to discriminate between barcodes, as it is more sensitive to Nanopore errors (described below) but also more computationally expensive.

#### Edit distance

The edit distance is defined as:

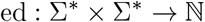

which returns the minimum number of insertions, deletions, and substitutions required to transform one string into another. Given strings *x* and *y*, we denote their distance as *d*:= ed(*x, y*).

#### Subsequence barcode scoring scheme

In Nanopore sequencing, errors often appear as stretches of nucleotides that are incorrect or missing, typically caused by slippage or stalling of DNA in the pore [41]. A single error stretch, e.g. TTTT, can already introduce four edits in an otherwise perfect alignment. In contrast, observing four edits scattered across an entire barcode is unlikely to result from such localized slippage or stalling errors (for an example see Figure 4).

To capture this distinction, we define a scoring scheme on top of the CIGAR string, *C*, of an edit-distance-based alignment. Specifically, we adapt subsequence scoring from Lodhi et al. [19] to operate on the CIGAR representation^3^. From the CIGAR string we can extract all query positions that matched (i.e. no substitutions, insertions, or deletions) in *P:*

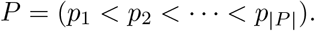

Then, given a subsequence length *k ≥* 1 and a decay parameter *λ* ∈ (0,1], we compute a score *S*_*k*_ that is large when the alignment contains many ordered groups of *k* matches that are tightly packed, and small when such groups are rare or interleaved with errors. We count every increasing *k*-tuple of match positions, weighting each group by an exponential penalty based on its span. Smaller values *λ* ≪ 1 penalize wide spacing more strongly, while *λ* ≈ 1 treats spacing more uniformly.

Formally, for *k* ≥ 1 and *λ* > 0, the score is

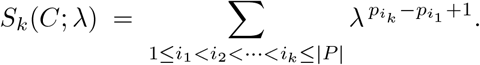

If |*P*| < *k*, then *S*_*k*_ = 0. For examples see Section A (Additional file 1).

### 5.3 Demultiplexing

Barbell has four main steps: `annotate`, `inspect`, `filter`, and `trim`.

#### Annotate

The `annotate` step refers to locating and scoring barcodes in the reads. In this manuscript we focus on rapid barcoding, but all Nanopore kits are supported (e.g. `SQK-RBK114.96` and `SQK-NBD114.96`). The algorithm is described in Algorithm 1. In short, the user supplies a Fasta file (or multiple Fasta files) containing the tag sequences, from which `Barbell` derives *τ*_*N*_ and *β*. `Barbell` then locates *τ*_*N*_ in the reads, extracts the masked region, and compares it to each barcode in *β*. Whether a barcode matches is based on the subsequence score. By default, the score for *b* should be ≥20% of the perfect score, and the difference between the top two should be ≥10%. If a barcode is found, this is reported as Ftag where the F denotes front, otherwise the flank is reported as Fflank. In case of dual-end barcodes, the user can provide an additional Fasta file with an Rtag (R for rear), of which the incomplete Rtag is reported as Rflank. For all its searches, `Barbell` uses `Sassy` with a default overhang penalty *α* = 0.5, that halves the edit cost for bases that align beyond the read boundary. This makes it possible to recover truncated tags that terminate at read ends.

#### Inspect

The `annotate` step results in annotations for each of the reads. To provide a comprehensive overview of the patterns in the data, `Barbell` groups reads into human-readable patterns. These same pattern representations are used in the `filter` step. Each tag has the form of <type> [<ori>,<label>, <pos>, <cutdirection>] where

- `<type>` Tag class, e.g. Ftag, Rtag, Fflank, Rflank.
- `<ori>` Strand orientation: `fw` or `rv`.
- `<label>` Barcode label of the tag, derived from the FASTA header. In `inspect`, `Barbell` focuses on the locations of the tags in the reads, and does not report the barcode labels (e.g., BC01, Section 2.5). In `filter`, the user can filter explicitly based on the barcode label for each tag, using the following options:
  – * — any barcode label.
  – `BC01` — only *tags* with barcode label equal to `BC01`
  – ≈`experimentl` — only *tags* with barcode labels containing the substring `experiment1`, e.g., `BC01_experimentl` or `BC02_experimentl` but not `BC03_experiment2`
  – `?` (e.g., `?l`) — defines a wildcard grouping. The same number enforces equality across tags that use it. For example, Ftag[…,? 1,…]__Rtag[..?l,…] matches reads where the Ftag and Rtag share the same barcode label (e.g., BC01-BC01), but not reads where the labels differ (e.g., BC01–BC02).
- `<pos>` Location specifier, e.g.:
  – @left(0..250) — barcode alignment starts within the first 250 bases of the read
  – @right(0..250) — barcode alignment ends within the last 250 bases of the read
  – @prev_left(0..250) — barcode starts within 250 bases right of the previous tag
- `<cut direction>` Trim direction used in the filter step, e.g.:
  – > > — keep sequence after the tag
  – < < — keep sequence before the tag

As example, Ftag[fw,*,@left(0..250)] includes all reads that have a forward barcode in the first 250 bases, regardless of its exact label (*). These patterns can be more complex as shown in Figure 6.

**Fig. 6:**
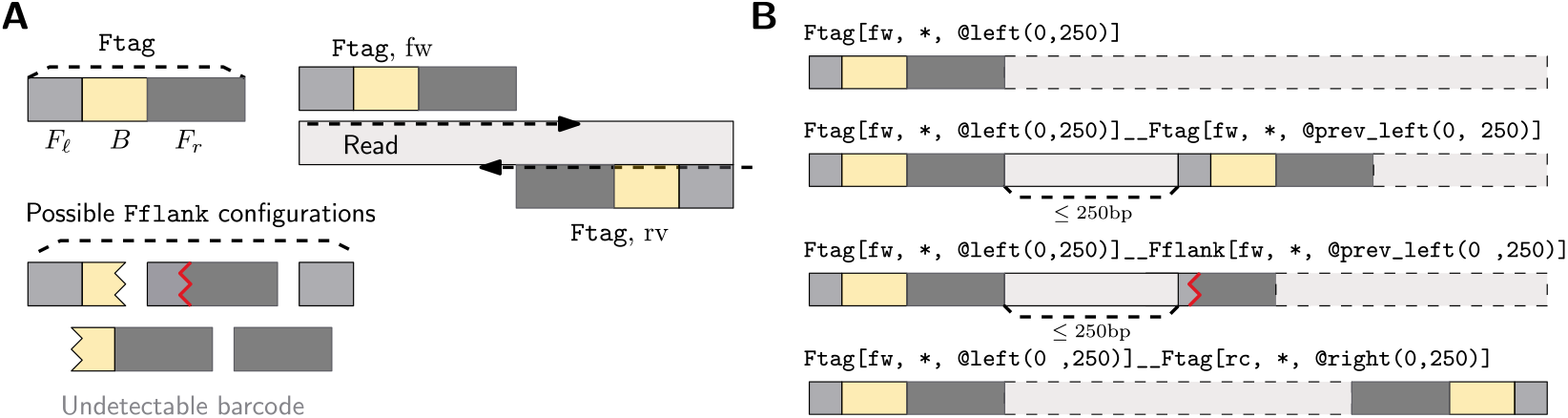
Example patterns observed in rapid barcoding data. **(A)** shows that an Ftag consists of the left flank (*F*_*ℓ*_), the barcode (*B*), and the right flank (*F*_*r*_). If *B* is undetectable based on the scoring scheme (e.g. absent or bad score) we report it as Fflank. If an Ftag, or Fflank matches the user provided sequence it is reported as `fw`, if it matches in reverse complement as `rc`. In **(B)** there are several examples of tag patterns observed in rapid barcoding data. The dashed borders (--) indicate the part of the read retained after trimming when using the `Barbell` rapid barcoding maximize preset. By default, a “grouping” of 250 bp is used, as tags are generally shorter than this, however, this value can be modified as a parameter in `inspect`.

#### Filter

The `filter` step lets a user extract the read annotations from `annotate` that match specific patterns. These can be directly copied from the overview reported by the `inspect` step, or manually tuned (e.g. only allowing a specific label). In the `filter` step, the user may also specify where reads should be cut to produce the desired trimmed read section. For example Ftag[fw,*,@left(0..250),>>] to trim off the Ftag on the left side and keep the section to the right (note the >>). This trim information is stored in the filtered annotation file.

#### Pattern ambiguity

We note that `inspect` and `filter` serve different purposes. `inspect` shows all patterns detected in the reads, whereas `filter` can be used to extract a subset of reads matching a certain pattern. Here, patterns are not necessarily unambiguous. For example a *tag* may match both @left(0..250) and @right(0..250) in the case of very short reads. Similarly, a *tag* might be close to the previous tag (@prev_left(0..250)) and the right end (@right(0..250)). `inspect` always prioritizes grouping based on @prev_left(i..j) over @right(i..j).

#### Trim

The `trim` step uses the `filter` results to trim the reads. For example, Ftag[fw,*,@left(0..250), >>] will retain the read section after the tag, and Ftag [fw,*,@right(0..250),<<] will retain the read section before the tag. Tags can be combined for dual-end barcoded reads, for example Ftag[fw,*,@left(0..250),>>]_ _Rtag[<<,rc,*,@right(0..250)] which trims both ends extracting the region between both barcodes.

### 5.4 Rapid barcoding patterns

For rapid barcoding we will consider the following two patterns safe:

1. Ftag[fw,*,@left(0..250),>>]
2. Ftag[fw,?1,@left(0..250)]_ _Ftag[fw,?1,@prev_left(0..250),>>]

The first pattern is the ideal pattern, with just a single left tag. The second pattern covers the second most common pattern (later in Results), where two barcodes are ligated for which we take the label from the inner tag (the one with >>), although we do enforce that both barcodes are the same using the ?l wildcard.

In case we want to maximize matches potentially at the expense of accuracy we add the following patterns:

1. Ftag[fw,*,@left(0..250)]_ _Ftag[fw,*,@prev_left(0,.250),>>]
2. Ftag[fw,*,@left(0..250),>>]_ _Ftag[<<,fw,*,@right(0..250)]
3. Ftag[fw,*,@left(0..250)]_ _Ftag[fw,*,@prev_left(0,.250),> >]_ _Ftag[< <,fw,*, @right(0..250)]

Here we always determine the sample based on the inner barcode, but are more flexible allowing additional barcodes to be present. Using maximize patterns will give most yield and should be used for tasks such as assembly, however for diagnostics and quantification, where false positives may affect the outcome, it might be better to use just the safe patterns.

### 5.5 Cut offs

A key step in demultiplexing is setting the thresholds that decide whether a region matching the flank *τ*_*N*_ and barcode *B* count as a match. Like other tools, `Barbell` uses edit distance to locate *τ*_*N*_, however we use a subsequence scoring scheme for the barcode region.

#### Edit distance cut-off

The expected edit distance between two random strings is on average 51% of their length, and can range between 36% and 63% [42]. In rapid barcoding, the flank *τ*_*N*_ has length |*τ*_*N*_| = 90, and the *N*_|*B*|_ mask of 24 N characters matches anything of the same length. Therefore, we define the effective flank length as

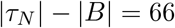

Based on the theoretical lower bound, we expect approximately

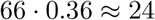

errors when matching against a random string. Because these theoretical values are derived from simulations of long strings, we fitted a lower bound through the edit distances of shorter strings (see Figure S1; Additional file 1 for details), yielding the formula

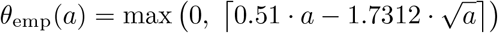

Here, *θ* represents the maximum number of edits, and the subscript “emp” indicates that it is an empirically fitted value based on our simulation.

This formula adjusts the theoretical 51% error rate downward for shorter sequences: the 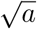 term grows sub-linearly with sequence length, imposing a stronger penalty on shorter sequences, which gradually diminishes for longer sequences.

Applying this to our effective flank length, we obtain

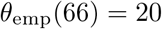

errors. We also verified the difference between a cut-off of 20, from the empirical data, versus 24 from Rosenfeld [42] and found that allowing 24 edits leads to some false positive flank matches within reads. `Barbell` implements the empirically derived formula that automatically sets the edit-distance cut-off according to the length of the user-provided tags. We report the automatic cut-off to the user such that it can be manually tuned when preferring more or less strict matching (Section 2.5).

#### Scoring barcodes

Both `Dorado` and `Flexiplex` use edit distance logic to identify the flanks and then the barcode within. `Flexiplex` sets a maximum edit distance (we used 6 in this manuscript), and if two barcodes have the same cost, none is returned. `Dorado` uses a more sophisticated heuristic where barcodes are only searched at expected locations (≤ 180 bases from the end), allowing up to 9 edits for the top hit, and being at least 3 edits from the second-best hit. If the top barcode has more than 9 edits, it should be 6 edits apart from the second barcode. This scoring is slightly more complex in reality as also flank scores are incorporated^4^. As described above (Section 5.2), we use a subsequence scoring scheme to score the barcode region. As the score, *S*, depends on *k* and *λ* setting a cut-off is not straightforward. To make this intuitive we first calculate a perfect score based on |*C*| matches. In case of barcodes that would be a CIGAR of 24 match operations (“24=“). Then the user can specify *S*_min_ and *S*_diff_ which are the percentage of the perfect score required to be considered a match, and the absolute percentage difference between the top two matches.

### 5.6 DNA isolation and sequencing

66 unidentified bacterial and fungal isolates were selected for Nanopore sequencing for diagnostic purposes (Table S1; Additional file 1). Briefly, genomic DNA was isolated using the DNeasy Ultra Clean Microbial kit (Qiagen, Venlo, the Netherlands). Nanopore sequencing was performed according to the rapid barcoding protocol RBK96.114 on an R10.4.1 flow cell with MinION (Oxford Nanopore, Oxford, UK). Bases were called using super accurate basecalling using MinKNOW v24.11.10.

### 5.7 Sequence searching and assemblies

For the analyses, we used `Sassy` [24] to search based on edit distance, always using an overhang of *α =* 0.5 (`-a 0.5`) to find matches crossing read boundaries and IUPAC alphabet to handle ambiguous bases (`--alphabet iupac`). To identify rapid barcoding contamination, we require that—aside from the flanks—a barcode is detected within ≤4 edits. Rapid barcoding kits use the *mu* transposase for barcode and adapter attachment. Since the *mu* transposase is naturally encoded by the *mu* phage, which infects *Enterobacteriaceae*, searching for just the flank could produce false positive matches—cases where the match reflects the presence of *mu* phage rather than true contamination. Annotation of our genomes showed two *Enterobacteriaceae* species in our own dataset. Moreover, when we later search databases these include many *Enterobacteriaceae* species.

As noted previously (Section 5.2), a typical rapid barcode flank consists of a left flank (*F*_*ℓ*_), a barcode (B), and a right flank (*F*_*r*_). In experimental data, however, we observed reads with two barcodes on the left side, following this concatenation configuration:

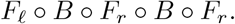

In this case, the second barcode entirely lacks its left flank and instead appears directly adjacent to the right flank of the preceding barcode region. We refer to such structures as fusions, and we searched for them in our datasets using the pattern GTTTTT CGTGCGCCGCTTCA<barcode_seq>GTTTTCGCATTTATCGTGAAACG. To detect fusions, we used MMseqs2 [43] using the following parameters: `-search-type 3, -max-seqs 5000000`, `-max-seq-len 200000`. We initially used `MMseqs2` instead of `Sassy`, since it was unclear whether fusion events would appear primarily as semi-global matches or also as shorter local sub-matches.

Typically, Filtlong [44] is used to discard the worst 10% of reads prior to assembly. Since this depends on how well the demultiplexer has already removed low-quality reads, we instead applied absolute thresholds. Reads were filtered with Filtlong (`v0.2.1`) keeping those ≥ 1000 bp (`-min_length 1000`) and with mean quality ≥ 15 (`-min_mean_q 15`). To assemble the genomes we used `Flye` (`v2.9.6-b1802`) [45] in -ont-hq mode with 5 polishing iterations (`-i 5`), followed by a final polishing using `Medaka` [46]. To map sequences to assemblies we used minimap2 (`v2.28-r1209`) [47], in `map-ont` mode (default parameters). To compare assemblies, we first extracted the contigs using Samtools [48], then mapped these to each other using Minimap2 (`map-ont`) [47], followed by graph induction using seqwish[49] and visualized using Bandage[50]

### 5.8 Tool comparisons

We observed that `Dorado` outputs untrimmed reads when a read consists entirely of barcode sequences. This is detectable as reads in the trimmed output file having the same length as in the original Fastq. While this behavior appears intentional^5^, it may be counterintuitive to users, who generally expect reads consisting solely of barcodes to be removed as implemented in `Flexiplex` and `Barbell`. In all analyses, we excluded 52,778 untrimmed reads outputted by `Dorado`; including these reads would increase the level of contamination. `Flexiplex` was designed for RNA-seq, and can report multiple barcodes per read by splitting the read. For our analyses, we retained only the longest fragment and associated barcode.

### 5.9 Taxonomic annotation

We used `Centrifuger` [51] with the RefSeq database [52] and Genome Taxonomy (GTDB) database (`r226`) [53]. As the GTDB alone does not include fungal sequences, we used the pre-pruilt GTDB + fungi database provided by `Centrifuger`. To link taxonomy identifiers to taxonomic lineages we use ete3 [54].

## Acknowledgements

We thank Torsten Schubert and Swapnil Doijad from the Viral Ecology and Omics (VEO) Group, Cluster of Excellence Balance of the Microverse, for providing Nanopore chemistry context and noticing the issue with `Dorado`’s trimming, respectively.

## Declarations

### 5.10 Ethics approval and consent to participate

Not applicable

### 5.11 Consent for publication

Not applicable

### 5.12 Availability of data and materials

Generated reads can be found under BioProject `PRJEB100828`. All code and steps to reproduce the results in this manuscripts can be found at GitHub (https://github.com/rickbeeloo/barbell-evals) all public data search results can be found at Zenodo (https://doi.org/10.5281/zenodo.17396505)

### 5.15 Author contributions statement

RB implemented the tool, conducted tests, and wrote the manuscript. RGK helped with the search algorithm (sassy), and provided valuable insights for the tool’s implementation. BED and AZ provided guidance for the experiments and analyses. XJ and AZ tested barbell on several datasets. LvIJ, EB and MBS sequenced isolates. All authors provided feedback on the manuscript.

## 5.17 Footnotes

^1^See https://github.com/nanoporetech/dorado/issues/626 for a discussion on several of these issues.

^2^ Filtered match results can be found at https://zenodo.org/records/17396505

^3^ For the implementation see our crate https://github.com/rickbeeloo/cigar-lodhi-rs.git

^4^ As there is no paper for `Dorado` please see the Dorado GitHub files `barcode_kits.h` and `BarcodeClassifier.cpp`.

^5^ In the demultiplexing code at https://github.com/nanoporetech/dorado/blob/release-v0.7/dorado/demux/Trimmer.cpp#L120-L125>, they specifically mention that trimming is skipped when the entire read consists of barcode sequence.

---

**Fig. S1;.**
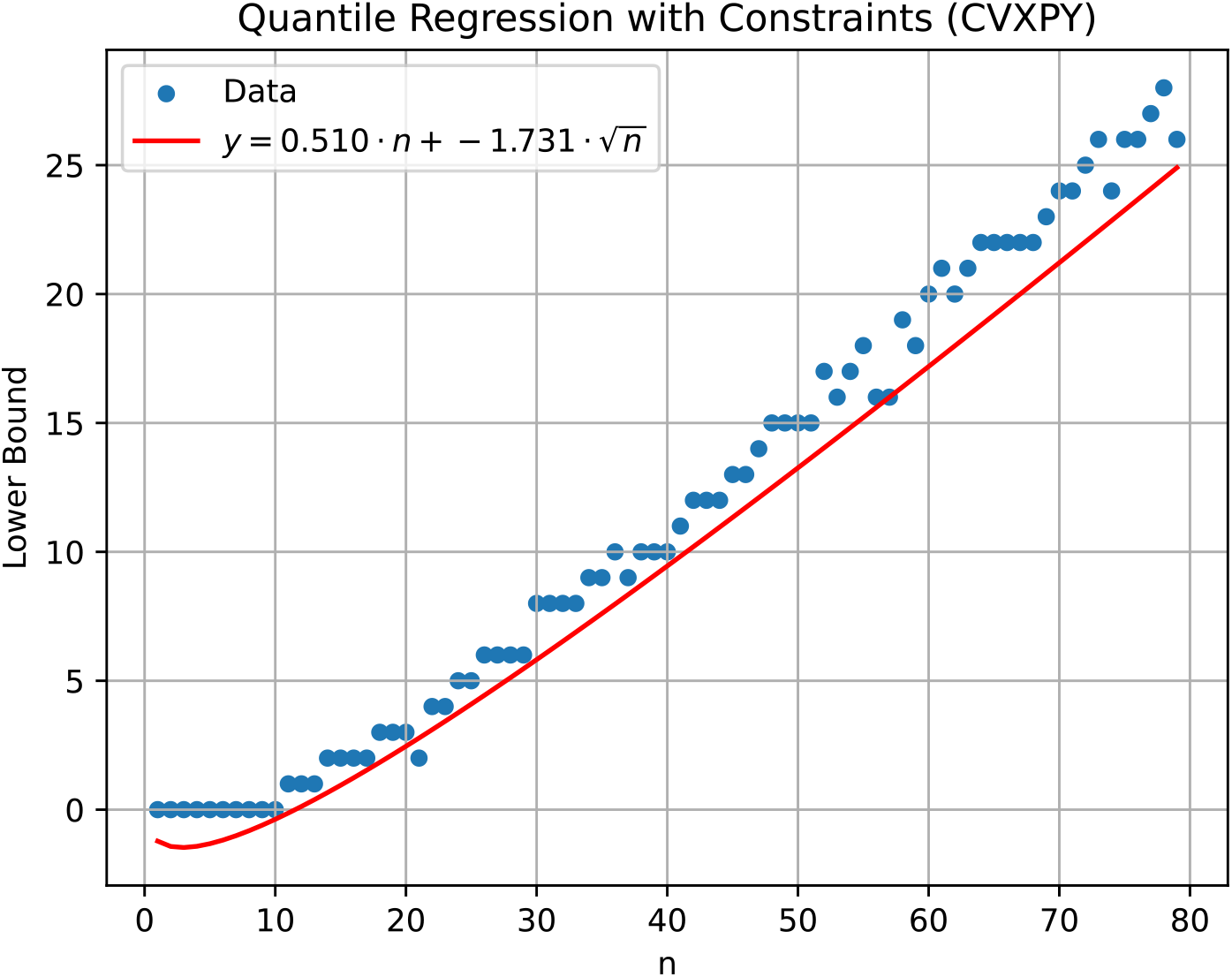
Additional file 1: Lower bound edit distance fit. For each length (x-axis) we performed 1 million random DNA versus random DNA comparisons, and plot the lowest observed edit distance (lower bound, y-axis). We then fitted a line through the lower 1% quantile, resulting in 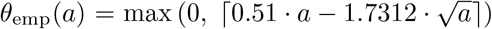 where we added a constraint to not allow negative values. This was solved uisng the Python package `cvxpy`.

## Appendix A Full calculation of CIGAR examples

We illustrate the score for *k* = 3 with decay *λ* on two CIGAR strings. Given match positions pos = (*p*_1_ < …< *p*_|_*C*|). the score is

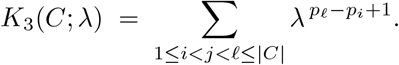

There is exactly one triple when |*C*| = 3, namely (*i, j, ℓ*) = (1,2,3), so 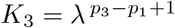.

**Fig. S2;.**
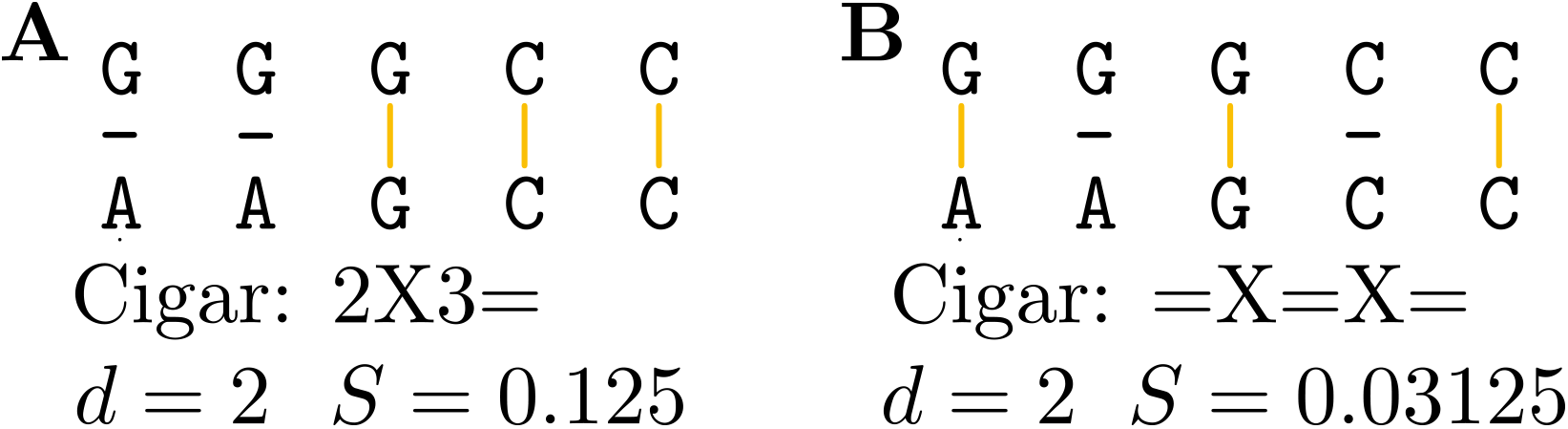
Additional file 1: Example of edit distance and subsequence scoring. Both alignments here have the same edit distance of 2, but in **(A)** the matches and errors are continguous, whereas in **(B)** the matches and errors are interleaved. Considering Nanopore errors arising from slippage and stalling of DNA in the pore, the alignment in (**A**) is more likely to be correct which is captured by the subsequence scoring (*S*).

### Example 1

`Sub Sub Match Match Match`

(Figure S2; Additional file 1A) Advancing the alignment index by each operation yields match positions POS = (2, 3, 4). The only 3-subsequence is (2, 3, 4) with inclusive span 4 – 2 + 1 = 3. hence

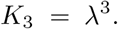

For 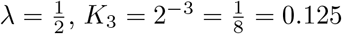.

### Example 2

`Match Sub Match Sub Match`

(Figure S2; Additional file 1B) Match positions are POS = (0,2,4). The only 3-subsequence is (0, 2, 4) with inclusive span 4 − 0 + 1 = 5, hence

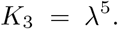

For 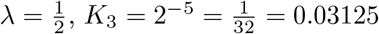.

### Interpretation

Both examples contain exactly one ordered triple of matches; the difference is the spacing between the first and last matches. The second CIGAR has larger gaps (due to substitutions), increasing the span and thus down-weighting the contribution more strongly via the _*λ*_^span^ factor.

## Appendix B Pore signal examples

To study the pore signal we used the raw pod5 files and basecalled these using dorado superaccurate model emitting the move table (`--emit-moves`). We then converted the pod5 files to slow5 using blue-crab [55], and visuzlized the pore signals and basecalled reads using Squigualiser [56], see Figure S4; Additional file 1. In case of secondary structure formation we expected the pore signal intensity (y-axis) to increase drastically (e.g. double), however this was not the case.

**Fig. S3;.**
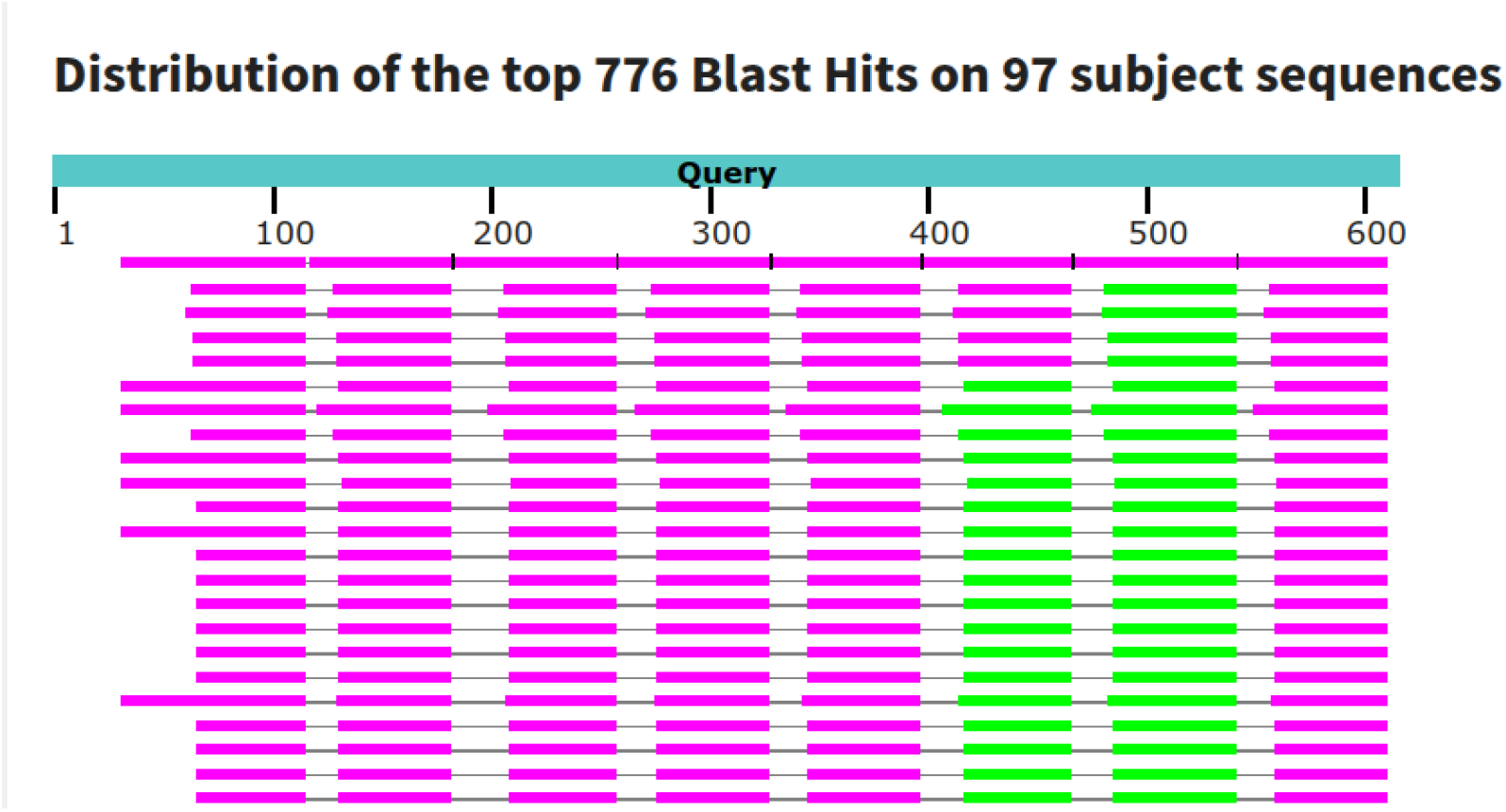
Additional file 1: Artefact read with 8 barcodes. This figure shows the BLAST output for read `52369018-4a3c-433b-881b-e46226500fb6` (611 bp) against all possible rapid barcode flanks and barcodes. The read consists entirely of barcode and flank sequences. `Barbell` detected 8× an Ftag in this read. The “Query” represents the read sequence. Each bar corresponds to a BLAST hit: pink bars indicate alignment scores of 80–200, and green bars 50–80. Because one region can match multiple barcode or flank sequences, matches appear underneath each other, with the highest-scoring ones shown on top. As expected, we observe eight distinct blocks (or “columns”), matching the number of Ftag’s detected by `Barbell`.

**Fig. S4;.**
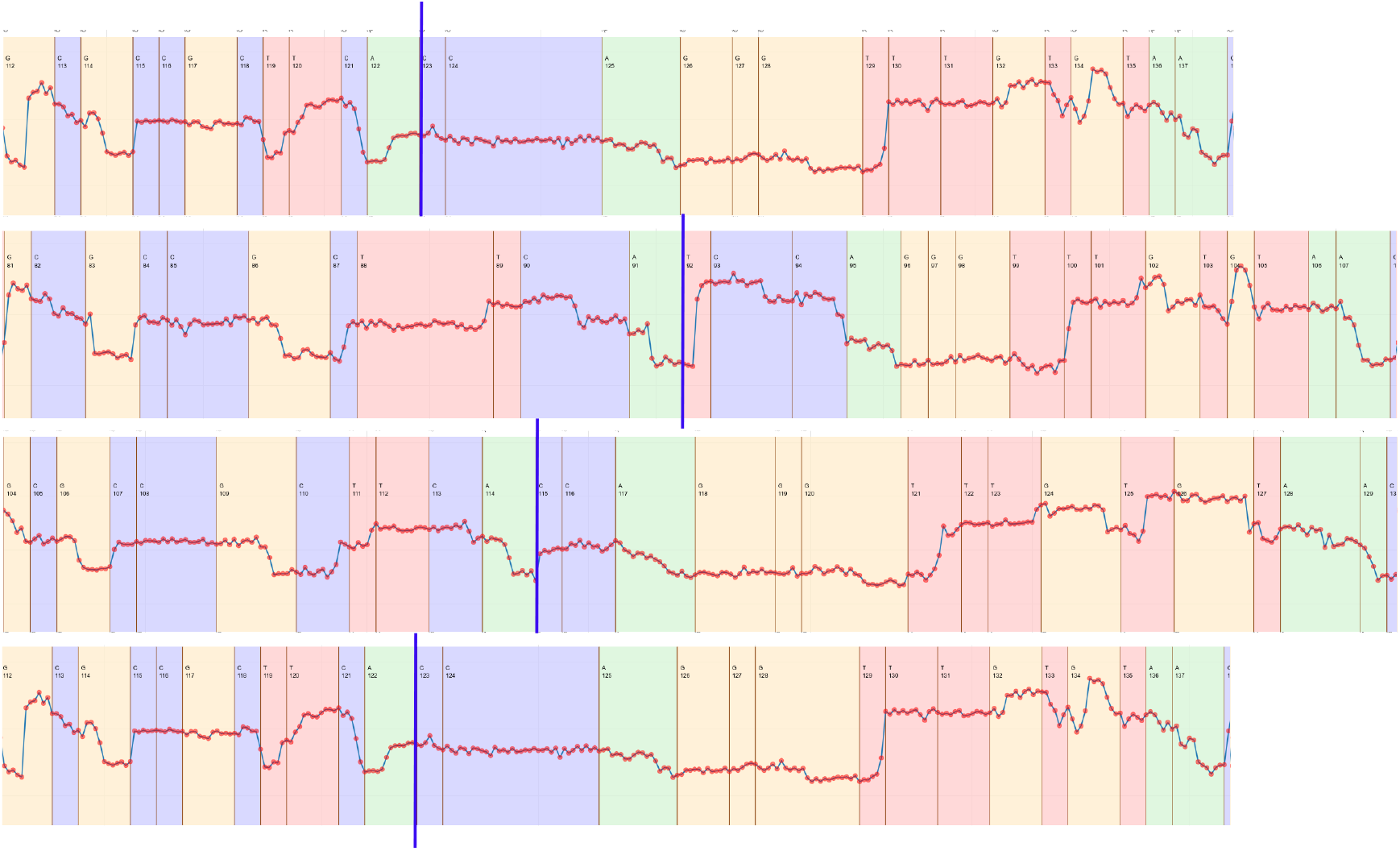
Additional file 1: Examples of the pore signal (line with red dots) for BC05 reads. The vertical blue line indicates the fusion point between the end of the right flank (`..GCTTCA`) and the beginning of the partial BC05 barocde (CTTGTCCAGGGTTTGTGTAACCTT). The colors indicate the basecalled bases, G=yellow, C=blue, T=red, A=green. We did not observe abnormally long stretches of signal without any basecalled bases.

**Table S1;.** Additional file 1: List of barcodes and their corresponding species assignments based on Centriguer (GTDB+Fungi), sorted by barcode. barcode02 and barcode03 are not included in the table due to insufficient reads for assembly, which prevented taxonomic annotation. We note that barcodes barcode44 and barcode46 are likely incorrectly annotated by `Centrifuger` and should be *Yersinia pseudotuberculosis* and *Brucella ceti*, respectively, based on more extensive analysis using an in-house pipeline

| Barcode | Species | Barcode | Species |
| --- | --- | --- | --- |
| BARCODE01 | *W11650 sp030535295* | BARCODE34 | *Cutibacterium acnes* |
| BARCODE04 | *Malassezia restricta* | BARCODE35 | *Fusobacterium russii* |
| BARCODE05 | *Castellaniella denitrificans* | BARCODE36 | *Mycobacterium abscessus* |
| BARCODE06 | *Psychrobacter sanguinis* | BARCODE37 | *Yamadazyma tenuis* |
| BARCODE07 | *JAUMYT01 sp030528525* | BARCODE38 | *Mycobacterium smegmatis* |
| BARCODE08 | *QD2021 sp036209505* | BARCODE40 | *Clostridium sp036643715* |
| BARCODE09 | *Exiguobacterium_A sp038006045* | BARCODE41 | *QD2021 sp036209505* |
| BARCODE10 | *Muribacter muris* | BARCODE42 | *Actinobacillus_C sp020026155* |
| BARCODE11 | *Acinetobacter terrestris* | BARCODE43 | *Mannheimia granulomatis* |
| BARCODE12 | *Pasteurella felis* | BARCODE44 | *Yersinia pestis* |
| BARCODE13 | *Prescottella sp032085135* | BARCODE45 | *Chelonobacter testudinis* |
| BARCODE14 | *Psychrobacter sanguinis* | BARCODE46 | *Brucella melitensis* |
| BARCODE15 | *Acinetobacter sp947627655* | BARCODE47 | *Actinobacillus_C sp020026155* |
| BARCODE16 | *Capnocytophaga catalasegens* | BARCODE48 | *Micrococcus luteus* |
| BARCODE18 | *Frederiksenia canicola* | BARCODE49 | *Saccharomyces cerevisiae* |
| BARCODE19 | *Granulicatella balaenopterae* | BARCODE50 | *Rodentibacter trehalosifermentans* |
| BARCODE20 | *Granulicatella balaenopterae* | BARCODE51 | *Actinomyces denticolens* |
| BARCODE22 | *Planococcus glaciei* | BARCODE52 | *Brevibacterium gallinarum* |
| BARCODE23 | *Fastidiosipila sp963510375* | BARCODE53 | *Streptococcus equi* |
| BARCODE24 | *Brachybacterium conglomeratum* | BARCODE54 | *Mannheimia haemolytica* |
| BARCODE25 | *Prescottella equi* | BARCODE56 | *Buchananella hordeovulneris* |
| BARCODE26 | *Prescottella equi* | BARCODE57 | *Staphylococcus simulans_B* |
| BARCODE27 | *Mannheimia haemolytica* | BARCODE59 | *QD2021 sp036209505* |
| BARCODE28 | *Carnobacterium maltaromaticum* | BARCODE60 | *Bisgaardia hudsonensis* |
| BARCODE29 | *Nicoletella semolina* | BARCODE61 | *Burkholderia thailandensis* |
| BARCODE30 | *Capnocytophaga stomatis* | BARCODE62 | *Intestinirhabdus alba* |
| BARCODE31 | *Gordonia sp016919385* | BARCODE63 | *Actinomyces denticolens* |
| BARCODE32 | *Berryella intestinalis* | BARCODE64 | *Streptococcus pasteurianus* |
| BARCODE33 | *Berryella intestinalis* | BARCODE65 | *Streptococcus gallolyticus* |
|  |  | BARCODE66 | *Streptococcus pasteurianus* |

### Algorithm 1: Pseudocode for Barbell’s annotate step.

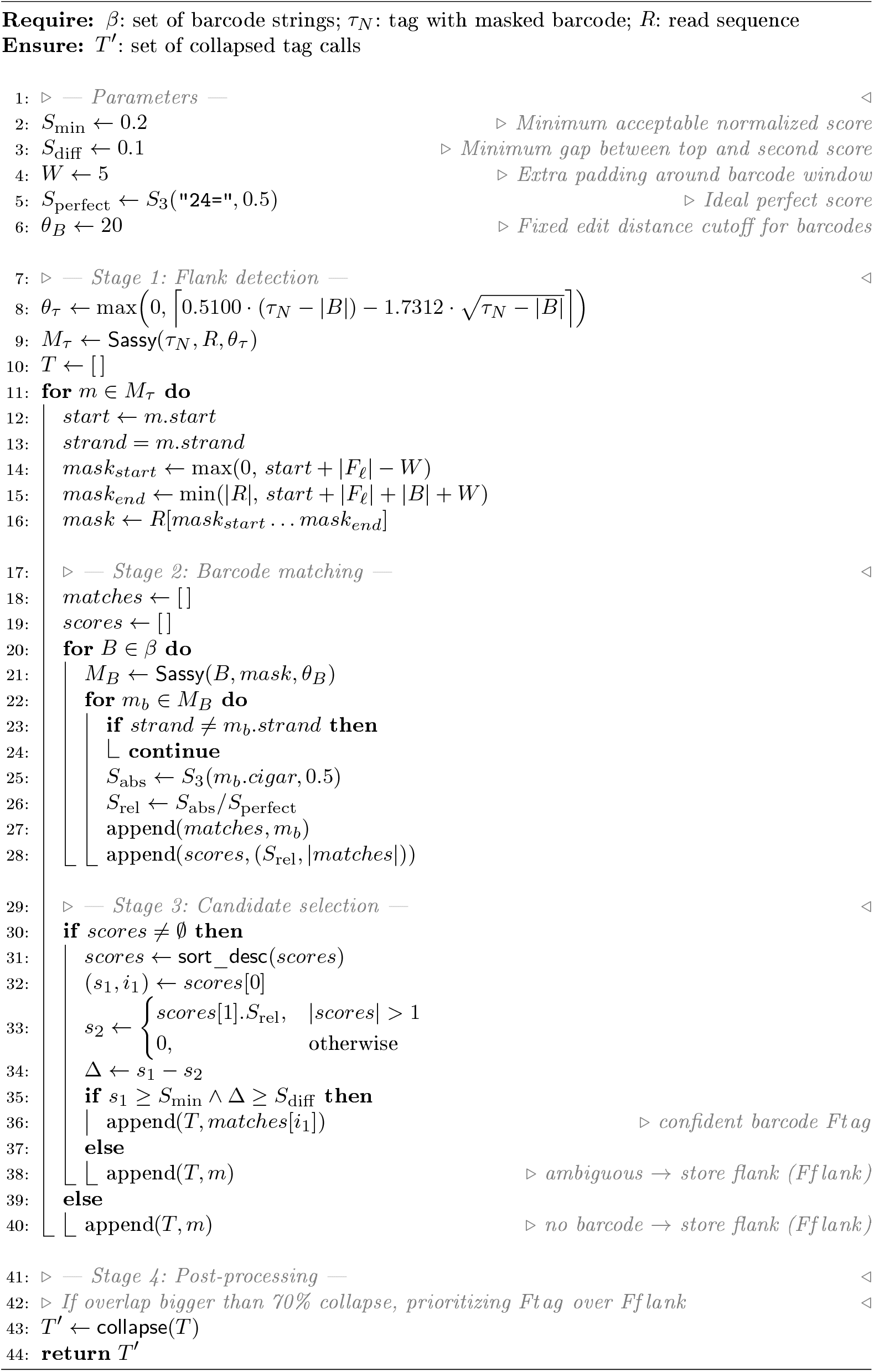

## References

[1] Kim, B.Y., Gellert, H.R., Church, S.H., Suvorov, A., Anderson, S.S., Barmina, O., Beskid, S.G., Comeault, A.A., Crown, K.N., Diamond, S.E., Dorus, S., Fujichika, T., Hemker, J.A., Hrcek, J., Kankare, M., Katoh, T., Magnacca, K.N., Martin, R.A., Matsunaga, T., Medeiros, M.J., Miller, D.E., Pitnick, S., Schiffer, M., Simoni, S., Steenwinkel, T.E., Syed, Z.A., Takahashi, A., Wei, K.H.-C., Yokoyama, T., Eisen, M.B., Kopp, A., Matute, D., Obbard, D.J., O’Grady, P.M., Price, D.K., Toda, M.J., Werner, T., Petrov, D.A.: Single-fly genome assemblies fill major phylogenomic gaps across the drosophilidae tree of life. PLOS Biology 22(7), 3002697 (2024) 10.1371/journal.pbio.3002697

[2] Karst, S.M., Ziels, R.M., Kirkegaard, R.H., Sørensen, E.A., McDonald, D., Zhu, Q., Knight, R., Albertsen, M.: High-accuracy long-read amplicon sequences using unique molecular identifiers with nanopore or pacbio sequencing. Nature Methods 18(2), 165–169 (2021) 10.1038/s41592-020-01041-y

[3] Wick, R.R., Judd, L.M., Holt, K.E.: Assembling the perfect bacterial genome using oxford nanopore and illumina sequencing. PLOS Computational Biology 19(3), 1010905 (2023) 10.1371/journal.pcbi.1010905

[4] Charalampous, T., Kay, G.L., Richardson, H., Aydin, A., Baldan, R., Jeanes, C., Rae, D., Grundy, S., Turner, D.J., Wain, J., Leggett, R.M., Livermore, D.M., O’Grady, J.: Nanopore metagenomics enables rapid clinical diagnosis of bacterial lower respiratory infection. Nature Biotechnology 37(7), 783–792 (2019) 10.1038/s41587-019-0156-5

[5] Li, Q., Zhao, X., Zhang, W., Wang, L., Wang, J., Xu, D., Mei, Z., Liu, Q., Du, S., Li, Z., Liang, X., Wang, X., Wei, H., Liu, P., Zou, J., Shen, H., Chen, A., Drmanac, S., Liu, J.S., Li, L., Jiang, H., Zhang, Y., Wang, J., Yang, H., Xu, X., Drmanac, R., Jiang, Y.: Reliable multiplex sequencing with rare index mis-assignment on dnb-based ngs platform. BMC Genomics 20(1) (2019) 10.1186/s12864-019-5569-5

[6] Vodák, D., Lorenz, S., Nakken, S., Aasheim, L.B., Holte, H., Bai, B., Myklebost, O., Meza-Zepeda, L.A., Hovig, E.: Sample-index misassignment impacts tumour exome sequencing. Scientific Reports 8(1) (2018) 10.1038/s41598-018-23563-4

[7] Sauvage, T., Cormier, A., Delphine, P.: A comparison of oxford nanopore library strategies for bacterial genomics. BMC Genomics 24(1) (2023) 10.1186/s12864-023-09729-z

[8] Xu, Y., Lewandowski, K., Lumley, S., Pullan, S., Vipond, R., Carroll, M., Foster, D., Matthews, P.C., Peto, T., Crook, D.: Detection of viral pathogens with multiplex nanopore minion sequencing: Be careful with cross-talk. Frontiers in Microbiology 9 (2018) 10.3389/fmicb.2018.02225

[9] Wu, X., Luo, H., Xu, F., Ge, C., Li, S., Deng, X., Wiedmann, M., Baker, R.C., Stevenson, A., Zhang, G., Tang, S.: Evaluation of salmonella serotype prediction with multiplex nanopore sequencing. Frontiers in Microbiology 12 (2021) 10.3389/fmicb.2021.637771

[10] Holzschuh, A., Lerch, A., Fakih, B.S., Aliy, S.M., Ali, M.H., Ali, M.A., Bruzzese, D.J., Yukich, J., Hetzel, M.W., Koepfli, C.: Using a mobile nanopore sequencing lab for end-to-end genomic surveillance of plasmodium falciparum: A feasibility study. PLOS Global Public Health 4(2), 0002743 (2024) 10.1371/journal.pgph.0002743

[11] Moeller, A.H., Dillard, B.A., Goldman, S.L., Real, M.V.F., Sprockett, D.D.: Removal of sequencing adapter contamination improves microbial genome databases. BMC Genomics 25(1) (2024) 10.1186/s12864-024-l0956-1

[12] Liu-Wei, W., Toorn, W., Bohn, P., Hölzer, M., Smyth, R.P., Kleist, M.: Sequencing accuracy and systematic errors of nanopore direct rna sequencing. BMC Genomics 25(1) (2024) 10.1186/s12864-024-10440-w

[13] Bonenfant, Q., Noé, L., Touzet, H.: Porechop_abi: discovering unknown adapters in oxford nanopore technology sequencing reads for downstream trimming. Bioinformatics Advances 3(1) (2022) 10.1093/bioadv/vbac085

[14] Šošić, M., Šikić, M.: Edlib: a C/C++ library for fast, exact sequence alignment using edit distance. Bioinformatics 33(9), 1394–1395 (2017)

[15] Logan, R., Fleischmann, Z., Annis, S., Wehe, A.W., Tilly, J.L., Woods, D.C., Khrapko, K.: 3gold: optimized levenshtein distance for clustering third-generation sequencing data. BMC Bioinformatics 23(1) (2022) 10.1186/s12859-022-04637-7

[16] Yu, T., Ren, Z., Gao, X., Li, G., Han, R.: Generating barcodes for nanopore sequencing data with pro. Fundamental Research 4(4), 785–794 (2024) 10.1016/j.fmre.2024.04.014

[17] Toorn, W., Bohn, P., Liu-Wei, W., Olguin-Nava, M., Gribling-Burrer, A.-S., Smyth, R.P., Kleist, M.: Demultiplexing and barcode-specific adaptive sampling for nanopore direct rna sequencing. Nature Communications 16(1) (2025) 10.1038/s41467-025-59102-9

[18] Pryszcz, L.P., Diensthuber, G., Llovera, L., Medina, R., Delgado-Tejedor, A., Cozzuto, L., Ponomarenko, J., Novoa, E.M.: Seqtagger, a rapid and accurate tool to demultiplex direct rna nanopore sequencing datasets (2024) 10.1101/2024.10.29.620808

[19] Lodhi, H., Saunders, C., Shawe-Taylor, J., Cristianini, N., Watkins, C.: Text classification using string kernels. Journal of machine learning research 2(Feb), 419–444 (2002)

[20] Li, H.-L., Pang, Y.-H., Liu, B.: Bioseq-blm: a platform for analyzing dna, rna and protein sequences based on biological language models. Nucleic Acids Research 49(22), 129–129 (2021) 10.1093/nar/gkab829

[21] Jia, X., Schubert, T., Beeloo, R., Litos, A., Doijad, S., Helvoort, P., Sperlea, T., Labrenz, M., Dutilh, B.E.: Bacterial community adaptation after freshwater and seawater coalescence (2025) 10.1101/2025.09.09.675091

[22] Sullivan, D.K., Pachter, L.: Flexible parsing, interpretation, and editing of technical sequences with splitcode. Bioinformatics 40(6) (2024) 10.1093/bioinformatics/btae33l

[23] Cheng, O., Ling, M.H., Wang, C., Wu, S., Ritchie, M.E., Göke, J., Amin, N., Davidson, N.M.: Flexiplex: a versatile demultiplexer and search tool for omics data. Bioinformatics 40(3) (2024) 10.1093/bioinformatics/btael02

[24] Beeloo, R., Groot Koerkamp, R.: Sassy: Searching short dna strings in the 2020s (2025) 10.1101/2025.07.22.666207

[25] Weinmaier, T., Conzemius, R., Bergman, Y., Lewis, S., Jacobs, E.B., Tamma, P.D., Materna, A., Weinberger, J., Beisken, S., Simner, P.J.: Validation and application of long-read whole-genome sequencing for antimicrobial resistance gene detection and antimicrobial susceptibility testing. Antimicrobial Agents and Chemotherapy 67(1) (2023) 10.1128/aac.01072-22

[26] Di Pilato, V., Bonaiuto, C., Morecchiato, F., Antonelli, A., Giani, T., Rossolini, G.M.: Next-generation diagnostics of bloodstream infections enabled by rapid whole-genome sequencing of bacterial cells purified from blood cultures. eBioMedicine 114, 105633 (2025) 10.1016/j.ebiom.2025.105633

[27] Tatusova, T., Ciufo, S., Fedorov, B., O’Neill, K., Tolstoy, L: Refseq microbial genomes database: new representation and annotation strategy. Nucleic Acids Research 42(Dl), 553–559 (2013) 10.1093/nar/gktl274

[28] Jørgensen, T.S., Mohite, O.S., Sterndorff, E.B., Alvarez-Arevalo, M., Blin, K., Booth, T.J., Charusanti, P., Faurdal, D., Hansen, T.O., Nuhamunada, M., Mourched, A.-S., Palsson, B.O., Weber, T.: A treasure trove of 1034 actinomycete genomes. Nucleic Acids Research 52(13), 7487–7503 (2024) 10.1093/nar/gkae523

[29] Lim, S.J., Natarajan, O., Keller, J., Dishaw, L.J., Furman, B.T., Breitbart, M.: Draft genome sequence of gracilimonas sp. strain bcb1 isolated from the gill tissue of the lucinid bivalve stewartia floridana in pinellas county, florida, usa. Microbiology Resource Announcements (2025) 10.1128/mra.00595-25

[30] Chklovski, A., Parks, D.H., Woodcroft, B.J., Tyson, G.W.: Checkm2: a rapid, scalable and accurate tool for assessing microbial genome quality using machine learning. Nature Methods 20(8), 1203–1212 (2023) 10.1038/s41592-023-01940-w

[31] Gould, A.L., Henderson, J.B.: Comparative genomics of symbiotic photobacterium using highly contiguous genome assemblies from long read sequences. Microbial Genomics 9(12) (2023) 10.1099/mgen.0.001161

[32] Matlock, W., Rodger, G., Pritchard, E., Colpus, M., Kapel, N., Barrett, L., Morgan, M., Oakley, S., Hopkins, K.L., Roohi, A., Karageorgopoulos, D., Avison, M.B., Walker, A.S., Lipworth, S., Stoesser, N.: E. coliphylogeny drives co-amoxiclav resistance through variable expression ofblatem-1 (2024) 10.1101/2024.08.12.607562

[33] Nondoli, H., Maghembe, R., Kidima, W., Makene, V., Ngadaya, E.: Complete genome sequences and multidrug resistance genotypes of nontuberculous mycobacteria isolates from the central tuberculosis reference laboratory muhimbili tanzania. Tanzania Journal of Health Research 25(2), 889–921 (2024) 10.4314/thrb.v25i2.15

[34] Li, X., Hu, H., Zhu, Y., Wang, T., Lu, Y., Wang, X., Peng, Z., Sun, M., Chen, H., Zheng, J., Tan, C.: Population structure and antibiotic resistance of swine extraintestinal pathogenic escherichia coli from china. Nature Communications 15(1) (2024) 10.1038/s41467-024-50268-2

[35] Rang, F.J., Kloosterman, W.P., Ridder, J.: From squiggle to basepair: computational approaches for improving nanopore sequencing read accuracy. Genome Biology 19(1) (2018) 10.1186/s13059-018-1462-9

[36] Greenfield, P., Duesing, K., Papanicolaou, A., Bauer, D.C.: Blue: correcting sequencing errors using consensus and context. Bioinformatics 30(19), 2723–2732 (2014) 10.1093/bioinformatics/btu368

[37] Liu, Y., Li, Y., Chen, E., Xu, J., Zhang, W., Zeng, X., Luo, X.: Repeat and haplotype aware error correction in nanopore sequencing reads with dechat. Communications Biology 7(1) (2024) 10.1038/s42003-024-07376-y

[38] Chen, Z., Ong, C.T., Nguyen, L.T., Lamb, H.J., González-Recio, O., Gutiérrez-Rivas, M., Meale, S.J., Ross, E.M.: Biases from oxford nanopore library preparation kits and their effects on microbiome and genome analysis. BMC Genomics 26(1) (2025) 10.1186/s12864-025-11649-z

[39] Salter, S.J., Cox, M.J., Turek, E.M., Calus, S.T., Cookson, W.O., Moffatt, M.F., Turner, P., Parkhill, J., Loman, N.J., Walker, A.W.: Reagent and laboratory contamination can critically impact sequence-based microbiome analyses. BMC Biology 12(1) (2014) 10.1186/s12915-014-0087-z

[40] Chen, S.: Ultrafast one-pass fastq data preprocessing, quality control, and deduplication using fastp. iMeta 2(2) (2023) 10.1002/imt2.107

[41] Delahaye, C., Nicolas, J.: Sequencing dna with nanopores: Troubles and biases. PLOS ONE 16(10), 0257521 (2021) 10.1371/journal.pone.0257521

[42] Rosenfeld, M.: Upper bounds on the average edit distance between two random strings (2024) 10.48550/ARXIV.2407.18113

[43] Steinegger, M., Söding, J.: Mmseqs2 enables sensitive protein sequence searching for the analysis of massive data sets. Nature Biotechnology 35(11), 1026–1028 (2017) 10.1038/nbt.3988

[44] Wick, R.R.: Filtlong: quality filtering tool for long reads. https://github.com/rrwick/Filtlong. Accessed: 2025-08-17 (2017)

[45] Kolmogorov, M., Yuan, J., Lin, Y., Pevzner, P.A.: Assembly of long, error-prone reads using repeat graphs. Nature Biotechnology 37(5), 540–546 (2019) 10.1038/s41587-019-0072-8

[46] Nanopore, O.: medaka. Accessed: 2025-10-02 (2020). https://github.com/nanoporetech/medaka

[47] Li, H.: Minimap2: pairwise alignment for nucleotide sequences. Bioinformatics 34(18), 3094–3100 (2018) 10.1093/bioinformatics/btyl91

[48] Li, H., Handsaker, B., Wysoker, A., Fennell, T., Ruan, J., Homer, N., Marth, G., Abecasis, G., Durbin, R.: The sequence alignment/map format and samtools. Bioinformatics 25(16), 2078–2079 (2009) 10.1093/bioinformatics/btp352

[49] Garrison, E., Guarracino, A.: Unbiased pangenome graphs. Bioinformatics 39(1) (2022) 10.1093/bioinformatics/btac743

[50] Wick, R.R., Schultz, M.B., Zobel, J., Holt, K.E.: Bandage: interactive visualization of de novo genome assemblies. Bioinformatics 31(20), 3350–3352 (2015) 10.1093/bioinformatics/btv383

[51] Song, L., Langmead, B.: Centrifuger: lossless compression of microbial genomes for efficient and accurate metagenomic sequence classification. Genome Biology 25(1) (2024) 10.1186/s13059-024-03244-4

[52] Pruitt, K.D., Tatusova, T., Maglott, D.R.: Ncbi reference sequences (refseq): a curated non-redundant sequence database of genomes, transcripts and proteins. Nucleic Acids Research 35(Database), 61–65 (2007) 10.1093/nar/gk1842

[53] Parks, D.H., Chuvochina, M., Rinke, C., Mussig, A.J., Chaumeil, P.-A., Hugenholtz, P.: Gtdb: an ongoing census of bacterial and archaeal diversity through a phylogenetically consistent, rank normalized and complete genome-based taxonomy. Nucleic Acids Research 50(D1), 785–794 (2021) 10.1093/nar/gkab776

[54] Huerta-Cepas, J., Serra, F., Bork, P.: Ete 3: Reconstruction, analysis, and visualization of phylogenomic data. Molecular Biology and Evolution 33(6), 1635–1638 (2016) 10.1093/molbev/msw046

[55] Gamaarachchi, H., Samarakoon, H., Jenner, S.P., Ferguson, J.M., Amos, T.G., Hammond, J.M., Saadat, H., Smith, M.A., Parameswaran, S., Deveson, I.W.: Fast nanopore sequencing data analysis with slow5. Nature Biotechnology 40(7), 1026–1029 (2022) 10.1038/s41587-021-01147-4

[56] Samarakoon, H., Liyanage, K., Ferguson, J.M., Parameswaran, S., Gamaarachchi, H., Deveson, I.W.: Interactive visualization of nanopore sequencing signal data with squigualiser. Bioinformatics 40(8) (2024) 10.1093/bioinformatics/btae5Ol

